# Hybrid adaptation is hampered by Haldane’s sieve

**DOI:** 10.1101/2023.12.15.571924

**Authors:** Carla Bautista, Isabelle Gagnon-Arsenault, Mariia Utrobina, Anna Fijarczyk, Devin P. Bendixsen, Rike Stelkens, Christian R. Landry

## Abstract

Hybrids between species exhibit plastic genomic architectures that foster phenotypic diversity. Their genomic instability also incurs costs, potentially limiting adaptation. When challenged to evolve in an environment containing a UV mimetic drug, yeast hybrids have reduced adaptation rates compared to parents. We hypothesized that this reduction could result from a faster accumulation of genomic changes, but we found no such association. Alternatively, we proposed that hybrids might lack access to adaptive mutations occurring in the parents, yet, we identified mutations in the same genes (*PDR1 and YRR1*), suggesting similar molecular adaptation mechanisms. However, mutations in these genes tended to be homozygous in the parents but heterozygous in the hybrids. We hypothesized that a lower rate of loss of heterozygosity (LOH) in hybrids could limit fitness gain. Using genome editing, we demonstrated that mutations display incomplete dominance, requiring homozygosity to show full impact and to circumvent Haldane’s sieve, which favors the fixation of dominant mutations. We used frozen ‘fossils’ to track genotype frequency dynamics and confirmed that LOH occurs at a slower pace in hybrids than in parents. Together, these findings show that Haldane’s sieve slows down adaptation in hybrids, revealing an intrinsic constraint of hybrid genomic architecture that can limit the role of hybridization in adaptive evolution.

## Introduction

Hybridization rapidly generates novel genotypes that can also lead to new and sometimes extreme phenotypes^1–5^. As a result, hybrids may thrive and often outcompete their parents^6–8^. Empirical and theoretical work has shown that hybridization can promote rapid evolution^2,3,9–15^, including during evolutionary rescue, species diversification, and adaptive radiations^2,12,16–19^. The generation of adaptive diversity through hybridization has long been successfully employed in biotechnology for fermentation^20–22^ and in agriculture for crop improvement^23–25^.

The combination of divergent genomes in the same organism can also lead to genomic instability^26,27^. From microorganisms^28,29^ to multicellular eukaryotes such as plants^30^ and vertebrates^31^, genomic instability has been frequently observed in hybrids. While the causes of instability are not always clear, evidence points to the alteration of the molecular pathways and components responsible for genome stability themselves. For instance, cell cycle checkpoints and DNA repair pathways observed to be altered in hybrids^32,33^ can lead to inaccurate chromosome segregation^34,35^, resulting in changes in ploidy^36–39^, aneuploidies^40–42^, and elevated mutation rates^43–46^.

In spite of these potential negative consequences, genomic instability can also paradoxically enhance F1 hybrid traits. One example is the restoration of meiotic recombination through genome homogenization, which contributes to rescuing hybrid fertility^47^. Similarly, whole-genome duplication can restore fertility in interspecific hybrids^48^. However, the enhancement of these hybrid traits is typically observed under stable laboratory conditions. Some environmental conditions and stresses could further enhance genome instability, so what appears as a feature that enhances adaptability in benign conditions could become a liability in extreme ones. Motivated by this question, in a previous study^49^, we produced F1 hybrids between the budding yeast *Saccharomyces cerevisiae* and *S. paradoxus* to measure their adaptive potential in an environment containing a drug that increases genomic instability by mimicking UV radiation^50,51^. These two species diverged 5-10 million years ago^52–56^, live in similar ecological niches in nature^57^, and carry signs for introgression in their mitochondrial and nuclear genomes with adaptive potential in some cases^47,58–61^. Replicated experimental evolution across 100 generations revealed that hybrids showed smaller fitness gains than their parents in conditions mimicking UV radiation^49^.

Here, we investigated the genomic and genetic basis of differential adaptive rates in hybrids by testing two non-mutually exclusive hypotheses: 1) Hybrid adaptation to UV radiation is hampered by genomic instability, and 2) Hybrids do not have access to the same adaptive mutations as the parental species. We used whole genome sequencing of 270 ancestral and evolved hybrid and parental genotypic backgrounds to: 1) Investigate major genomic changes in copy number such as ploidy changes, aneuploidies, or loss of heterozygosity (LOH), to determine if they are more prevalent in hybrid genomes, and 2) Identify potentially differential *de novo* mutations in hybrids compared to parents. We functionally validated seven putative adaptive mutations with site-directed mutagenesis in both *S. cerevisiae* and *S. paradoxus* haploid backgrounds and by CRISPR-Cas9 genome editing. We ultimately determined differences in the genetic architecture of adaptive mutations between hybrids and parents by tracking the allelic frequency over time through historical resurrection of frozen fossils. We found that none of our two starting hypotheses were supported. Rather, we found that adaptation to UV mimetic conditions often proceeds through two types of mutations, the second occurring more slowly in the hybrids, and thus slowing down their rate of adaptation.

## Results

### Hybrid and Parent Fitness in UV mimetic conditions

We previously evolved 90 populations (30 F1 hybrid replicates and 30 replicates of each of the parental *S. cerevisiae* and *S. paradoxus* genotypic backgrounds) for 100 generations in the presence of a DNA damaging agent, the UV mimetic drug 4-nitroquinoline 1-oxide (4-NQO), and in a control condition (rich medium) (*Fig. 1a*)^49^. We found that hybrids achieved a lower rate of adaptation compared to the parental genotypic backgrounds, with a lower average gain of fitness over the course of evolution^49^.

**Fig. 1.**
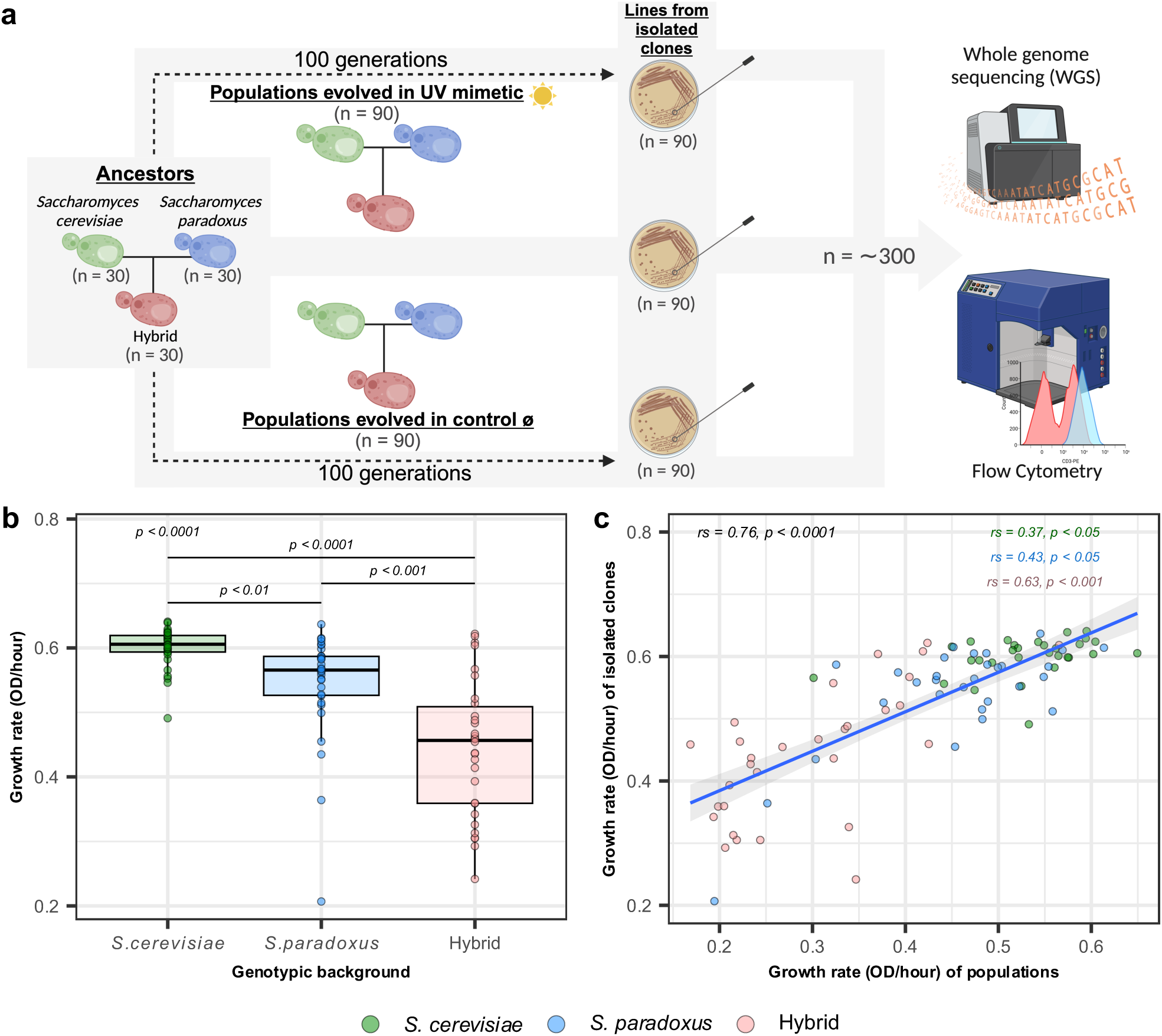
Hybrids show reduced adaptive potential in UV mimetic conditions. **a,** Experimental evolution in UV mimetic and control conditions was performed in hybrid and parents for about 100 generations^49^ (*n* = 30 replicated lines for each genotypic background: *S. cerevisiae*, *S. paradoxus* or hybrid). Lines from isolated clones derived from each population were analyzed by Whole Genome Sequencing (WGS) and flow cytometry (*n* = 300). **b,** Growth rate of lines from isolated clones derived from each population evolved in UV mimetic conditions (4 µM of 4-NQO) (*n* = 30 for each genotypic background: *S. cerevisiae*, *S. paradoxus* or hybrid). *p*-value for ANOVA (above) and Tukey post hoc pairwise *p*-values are shown. **c,** Growth rates of lines from isolated clones derived from each population correlate with growth rates of their populations of origin. Spearman’s rank correlation coefficients (rs) and associated *p*-values are shown (*n* = 30 for each genotypic background: *S. cerevisiae*, *S. paradoxus* or hybrid). Illustrations in **a** were created with BioRender.com.

To compare the number and type of genetic changes between evolved hybrid and parental populations, we measured DNA content by flow cytometry and sequenced the genomes of 300 isolated clones derived from the evolved populations and their ancestors (*Fig. 1a* and *Supplementary Table S1*) (average genome-wide coverage of 100X, *Supplementary Fig. 1*). In parallel, we measured the fitness of these isolated clones in control and UV mimetic conditions (*Extended Data* Fig. 1). We confirmed our previous finding that evolved hybrids showed significantly lower growth rate improvements than both parental replicated populations when evolved under UV mimetic conditions (*Fig. 1b,* all *p* < 0.001). The growth rates of isolated clones and that of their populations of origin were strongly correlated (*Fig. 1c*: *rs* = 0.76, *p* < 0.0001), confirming that the fitness of isolated clones is representative of the general fitness dynamics observed in the previous evolution experiment^49^. From now on, we will refer to lines when discussing observations on these isolated clones derived from their populations of origin.

### Major Genomic Changes in Copy Number Reveal Increased Genomic Instability in UV Mimetic Conditions

The majority of lines remained diploid during evolution. Ploidy changes were generally not more frequent under UV mimetic conditions (*Extended Data* Fig. 2*, Supplementary Fig. 2*). Likewise, we did not see a larger number of genomic changes in copy number in hybrids in UV mimetic compared to control conditions, suggesting that ploidy changes do not account for the observed reduction in hybrid adaptive potential. Conversely, evolution under UV mimetic conditions did result in a larger number of lines with aneuploidies in hybrid and both parental genotypic backgrounds when compared to control conditions (*Fig. 2a and 2b*), confirming that the UV mimetic treatment affected genome stability. However, the number of aneuploidies did not correlate with fitness changes observed in any group (*Supplementary Fig. 3*, *rs* = -0.18, *p* > 0.5; *rs* = -0.22, *p* > 0.5 and, *rs* = -0.27, *p* > 0.5; for *S. cerevisiae*, *S. paradoxus* and hybrid respectively). Hybrids also did not show a higher number of aneuploidies compared to the parents, making it unlikely to explain their reduced adaptability (*Fig. 2b*).

**Figure 2.**
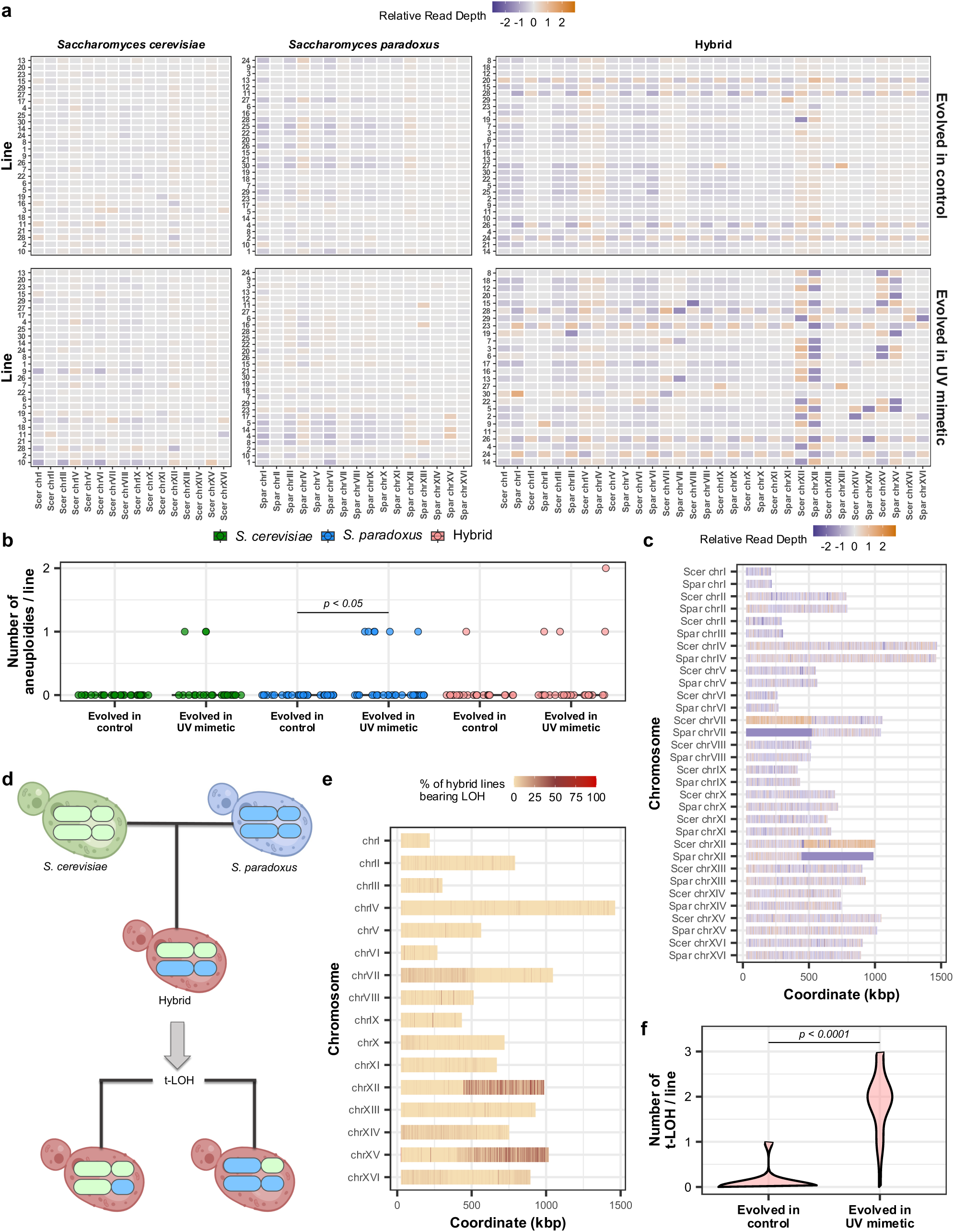
Genomic changes observed during experimental evolution. **a,** Relative read depth per chromosome and evolved line. Colored squares show the relative read depth between evolved and ancestor lines (log2 fold change). Values with an increase in 30% fold change represent gains in DNA content (gradient towards orange) and values with a decrease in 30% fold change represent losses in DNA content (gradient towards purple). Rows are individual genomes and columns are the chromosomes. Hybrid lines have two sets of chromosomes, since a concatenated hybrid genome was used for mapping. Homologous chromosomes are positioned side-by-side: *S. cerevisiae* (Scer) chromosomes on the left and *S. paradoxus* (Spar) on the right (See methods for details). Top panels show control lines and the bottom ones show lines evolved in UV mimetic conditions. Lines are ranked according to their fitness gain under UV mimetic conditions (with top lines indicating greater fitness). This same ranking order is preserved under control conditions, enabling a direct comparison between conditions (*n* = ∼30 lines per genotypic background and condition). **b,** Number of aneuploidies per line after evolution under control and UV mimetic conditions for the three genotypic backgrounds (*n* = ∼30 lines for each genotypic background and condition). Fisher’s Exact Test (within conditions for the same genotypic background) was performed. Only significant *p*-values are shown. **c,** Example of read depth variation across chromosomes for a single hybrid line (13) evolved in UV mimetic conditions, highlighting the detection of t-LOH through simultaneous increase and decrease in read depth (deviations of 30% from the genome-wide median read depth). **d,** Illustrative scheme of t-LOH produced in hybrid genomes. **e,** Percentage of hybrid lines carrying t-LOHs on each chromosome region (*n* = 24 hybrid lines). **f,** Number of t-LOHs per hybrid line after evolution under control and UV mimetic conditions. Wilcoxon test *p*-value is shown (*n* = 27 hybrid lines evolved in control and *n* = 24 hybrid lines evolved in UV mimetic). Illustrations in **d** were created with BioRender.com.

The analysis of the sequence depth of coverage revealed a particular type of alteration, for instance, in chromosomes XII and XV (*Fig. 2a*). Intriguingly, gains and losses occurred simultaneously in the homologous chromosomes of *S. cerevisiae* and *S. paradoxus,* i.e., when a hybrid lost a portion of the *S. cerevisiae* chromosome, it simultaneously gained a portion of the homologous *S. paradoxus* one. This prompted us to map the chromosome coverage to determine patterns in the distribution of species-specific chromosome gains and losses. The observed changes affected large regions of chromosomes (*Fig. 2c*). These patterns can be caused by the non-reciprocal exchange of homologous chromosomes in diploids during mitosis, resulting in loss of heterozygosity (LOH) (*Fig. 2d*)^62,63^. We identified LOH as regions where there were simultaneous increases and decreases in read depth in homologous chromosomes along a size threshold of 20 kb (*Extended Data* Fig. 3). We detected two types of LOH, interstitial LOHs (i-LOH), which often originate from gene conversion involving short exchanges, and terminal LOHs (t-LOH), typically resulting from mitotic crossovers that encompass larger regions^62,64^. We identified some large i-LOHs (for example in line 17 on chromosome XIV, *Extended Data* Fig. 3) but here we focused on t-LOHs, which were more frequent.

The frequency of t-LOH events in hybrid genomes was significantly positively correlated with chromosome size (*Extended Data* Fig. 4a*, rs* = 0.81, *p* < 0.05). The pattern of t-LOHs showed similarities across hybrid lines, with specific regions concentrated on chromosomes VII, XII, and XV (*Fig. 2e*). These regions have been previously identified to be susceptible for t-LOHs^65^. Certain t-LOHs were found to cluster around particular positions enriched with repetitive loci, such as the rDNA locus on chromosome XII^62,66^ or the *STE4* gene, another common t-LOH-target on chromosome XV^67^. We found a much larger number of hybrid lines with t-LOHs when evolved under UV mimetic conditions (96% of the lines) compared to control conditions (8% of the lines) (Fig. 2f, *p* < 0.0001). Only one line evolved under UV mimetic conditions did not show any t-LOH (4%).

Overall, our data indicates that t-LOHs are a specific outcome of evolution in UV mimetic conditions. This suggests that either DNA damage triggers t-LOH, thus enhancing its occurrence rate, or that t-LOH is particularly advantageous under these conditions and selected for. Advantageous LOHs have been associated with fitness increases in both lab and natural settings^65,68^. However, LOHs in themselves may not cause variation in the rates of adaptation among hybrid lines. We indeed did not find a significant correlation between the increase in fitness and the number of t-LOH events (*Extended Data* Fig. 4b, *rs* = 0.33, *p* > 0.05).

In summary, although we cannot examine t-LOHs in parental genomes due to their starting homozygosity, we found that aneuploidy frequencies are similar across parental species and hybrids, suggesting that overall genome instability is not specifically increased in hybrids. There is therefore no support for our initial hypothesis that increased instability hampers adaptation in hybrids.

### Parents and Hybrids have Parallel Access to Adaptive *De Novo* Mutations in the same Genes

Lower rates of adaptation to UV mimetic conditions may also be explained by hybrids not having access to the same adaptive mutations as the parental lines, as per our alternative hypothesis. For instance, some mutations could have strong genetic background-dependent effects. We investigated whether hybrids show parallel or distinct patterns in potentially adaptive single nucleotide polymorphisms (SNPs), focusing on missense variants, as their potential impacts are easier to interpret, and data is more robust in this category. We found a median of 20 missense variants per line in *S. cerevisiae*, 12 in *S. paradoxus*, and 42 in the hybrids (*Extended Data* Fig. 5) with a comparable trend in control conditions. A gene ontology (GO) analysis revealed differences in terms of the biological functions affected by mutations but also some similarities (*Supplementary Fig. 4*). Although non-significant, the most enriched GO terms for *S. cerevisiae* included double-strand break repair via sister chromatid exchange (GO:1990414), potentially aiding in coping with 4-NQO-induced DNA breaks, and the regulation of cell differentiation (GO:0045595). For *S. paradoxus*, the most enriched terms included ER-associated misfolded protein catabolic processes (GO:0071712) and endocytic vesicles (GO:0030139), potentially disrupting the invagination of extracellular substances, such as the 4-NQO drug (UV mimetic conditions). In the hybrid, the most enriched terms comprised trehalose metabolic process (GO:0005991) and ABC-type transporter activity (GO:0140359), comprising efflux pumps involved in expelling xenobiotic compounds, such as 4-NQO.

We found that mutations occurred in parallel in two specific genes across all three genotypic backgrounds (*Fig. 3a*), suggesting that the same molecular changes provide strong selective advantages in all three genotypic backgrounds. The most frequent parallel changes involved the pleiotropic drug response genes *PDR1* and *YRR1* (*Fig. 3a*). *PDR1* and *YRR1* encode zinc finger transcription factors regulating multidrug and stress responses^69–71^. Among targets, they modulate the expression of *PDR5*, a well-characterized yeast efflux pump that actively transports toxic compounds out of the cell^72,73^. Three *S. cerevisiae* lines, eight *S. paradoxus* lines, and eight hybrid lines had non-synonymous mutations in *PDR1*, while each genotypic background had five lines with *YRR1* mutations (*Fig. 3a*). This is a surprising level of parallelism, given that drug resistance mutations have been shown to be genotype-specific^74^ and that *S. cerevisiae* and *S. paradoxus* diverged 10 million years ago.

**Figure 3.**
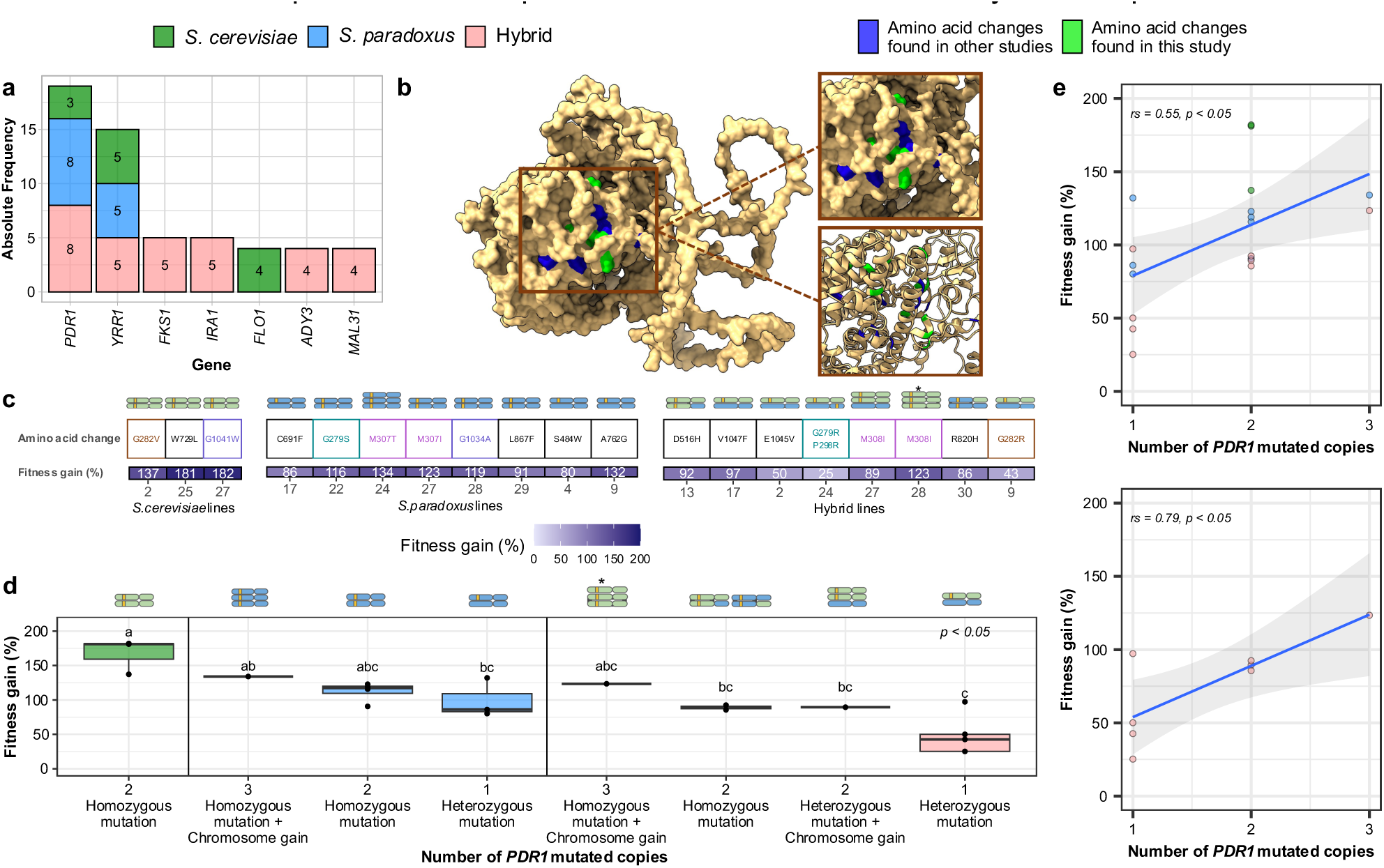
*PDR1* shows parallel adaptive changes among genotypic backgrounds. **a,** Absolute frequency of the most recurrent mutated genes in parents and hybrids. **b,** Pdr1p structure modeled with AlphaFold featuring amino acid changes identified in this study, alongside amino acid changes previously reported. All changes occur in the same regions. Cluster of amino acid changes is shown in the insets. **c,** Mutations and chromosomal changes occur together and impact fitness in the three genotypic backgrounds. A schematic of the chromosomal changes for each individual line is shown (*n* = 3 for *S. cerevisiae*, *n* = 8 for *S. paradoxus* and, *n* = 8 for hybrid). The number of *PDR1* mutated copies is shown in yellow. Identical amino acid changes are indicated by matching colors across the different genotypic backgrounds. **d,** Fitness gain (% change in growth rate between initial and final time points) as a function of the number of *PDR1* mutated copies (*n* = 3 for *S. cerevisiae* homozygous mutation, *n* = 1 for *S. paradoxus* homozygous mutation + chromosome gain*, n* = 4 for *S. paradoxus* homozygous mutation, *n* = 3 for *S. paradoxus* heterozygous mutation, *n* = 1 for hybrid homozygous mutation + chromosome gain, *n* = 2 for hybrid homozygous mutation, *n* = 1 hybrid heterozygous mutation + chromosome gain, and *n* = 4 for hybrid heterozygous mutation). *p*-value for Kruskal-Wallis test (above) and adjusted *p*-values after false discovery rate (FDR) multiple test correction (above each boxplot) are shown. The number of *PDR1* mutated copies is also shown in yellow. **e,** Fitness gain (% change in growth rate between initial and final time points) as a function of the number of *PDR1* mutated copies in the three genotypic backgrounds (top) or only in hybrid (bottom). Spearman’s rank coefficient (rs) and *p*-value are shown (*n* = 3 for *S. cerevisiae*, *n* = 8 for *S. paradoxus,* and *n* = 8 for hybrid).

From here on, we will focus on the analysis of *PDR1* since it was the most often mutated gene. Non-synonymous mutations in *PDR1* were not randomly scattered along the gene but instead occurred in particular clusters that overlapped among all three genotypic backgrounds (*Extended Data* Fig. 6). To identify the specific locations of the mutations, we analyzed the protein structure of Pdr1 (*Fig. 3b*). We found identical or similar substitutions as revealed in previous studies on *S. cerevisiae* exposed to different drugs, ethanol and antifungal molecules^75–79^(*Fig. 3b*). Localization on the protein structure revealed the presence of a cluster also found in other fungal species, for instance, in the pathogenic fungus *Nakaseomyces glabratus* (*Extended Data* Fig. 7a and 7b), for which antifungal resistance often arises from mutations in *PDR1*^80–86^. Parallelism even occurred at the level of amino acid changes across genotypic backgrounds (*Fig. 3c)*. For instance, a mutation at amino acid 308/307 (corresponding to the respective *S. cerevisiae* and *S. paradoxus* coordinates) independently occurred up to four times. This specific amino acid change has also been observed in other studies^76^, conferring resistance to the same UV mimetic drug we employed (4-NQO)^75^.

The identification of shared mutational hotspots and shared amino acid changes suggests a common adaptive landscape across genotypic backgrounds, indicating that specific regions within *PDR1* and *YRR1* harbor similar potential for adaptive mutations to occur in both hybrid and parents.

### Fitness increases with Copy Number of *PDR1* Adaptive Alleles

The occurrence of mutations in the *PDR1* gene across all three (hybrid and two parental) genotypic backgrounds raises an intriguing question: Why do these mutations not confer similar adaptive benefits to hybrids as they do to the parental species? We observed that some lines carrying *PDR1* mutations showed particularly high fitness gains under UV mimetic conditions. Specifically, lines 2, 25, and 27 for *S. cerevisiae*, lines 22, 24, 27, and 28 for *S. paradoxus*, and lines 13, 27, 28, and 30 for the hybrids show notable improvements (*Fig. 3c*). A closer genomic analysis revealed that all these lines share a common characteristic, namely the presence of multiple mutated copies of *PDR1* (*Fig. 3c*). Although mutations are expected to initially be heterozygous, we observed a diversity of genotypes at the *PDR1* locus. While some lines have undergone LOH that made the mutations homozygous, others show changes in ploidy resulting in an increased number of chromosomes harboring *PDR1* mutations (*Fig. 3c*). It is worth noting that an exception is observed in line 9 of *S. paradoxus* (*Fig. 3c*). While being heterozygous for the *PDR1* mutation, this line’s high fitness is likely due to an additional mutation in the *YRR1* transcription factor, resulting in enhanced copies of these two transcription factors simultaneously.

Fitness increased as a function of the number of *PDR1* mutated copies in all three genotypic backgrounds (*Fig. 3d* and *Fig. 3e* top panel *rs* = 0.55, *p* < 0.05). This effect was particularly strong in hybrids (Fig. 3e bottom panel, *rs* = 0.79, *p* < 0.05). The two lines with the largest observed fitness gain within their respective genotypic contexts (134% in *S. paradoxus* line 24 and 123% in hybrid line 28) each carried three copies of the *PDR1* mutant alleles (*Figure 3c and 3d*). Hybrid line 28 showed a complex pattern in which a whole-chromosome t-LOH was combined with an increase in ploidy (*Fig. 3c*, Supplementary Fig. 5 and Extended Data Fig. 3*).

The likelihood and rate of a mutation becoming fixed in a population is shaped by the strength of selection and the architecture of the trait under selection, including allelic dominance. Therefore, not only does an adaptive mutation need to occur, but it also needs to be in a proper genotype for the individuals to fully benefit from it. Mutations with greater dominance are more likely to become fixed, a principle referred to as Haldane’s sieve^87–89^. This poses a problem to non-obligate sexual species and in systems that only rarely reproduce sexually, like many unicellular microbes such as fungi. In asexual populations, recessive or incompletely dominant beneficial mutations can bypass Haldane’s sieve by achieving homozygosity through LOH ^90^, although completely recessive alleles would be invisible to selection and could thus be lost before an LOH occurs. The rate of LOH is not uniform among genotypes and it has been shown to be lower in heterozygous genotypes^91^. In hybrids, which are highly heterozygous, successful LOH rates are even more limited (Fig. 4a)^42,45^. Once a beneficial mutation occurs in the parental genomes, high rates of mitotic recombination can rapidly lead to LOH at the site where the mutation occurred and make a mutation homozygous. This would occur at a lower rate in hybrids, slowing down the rate of adaptation.

**Figure 4.**
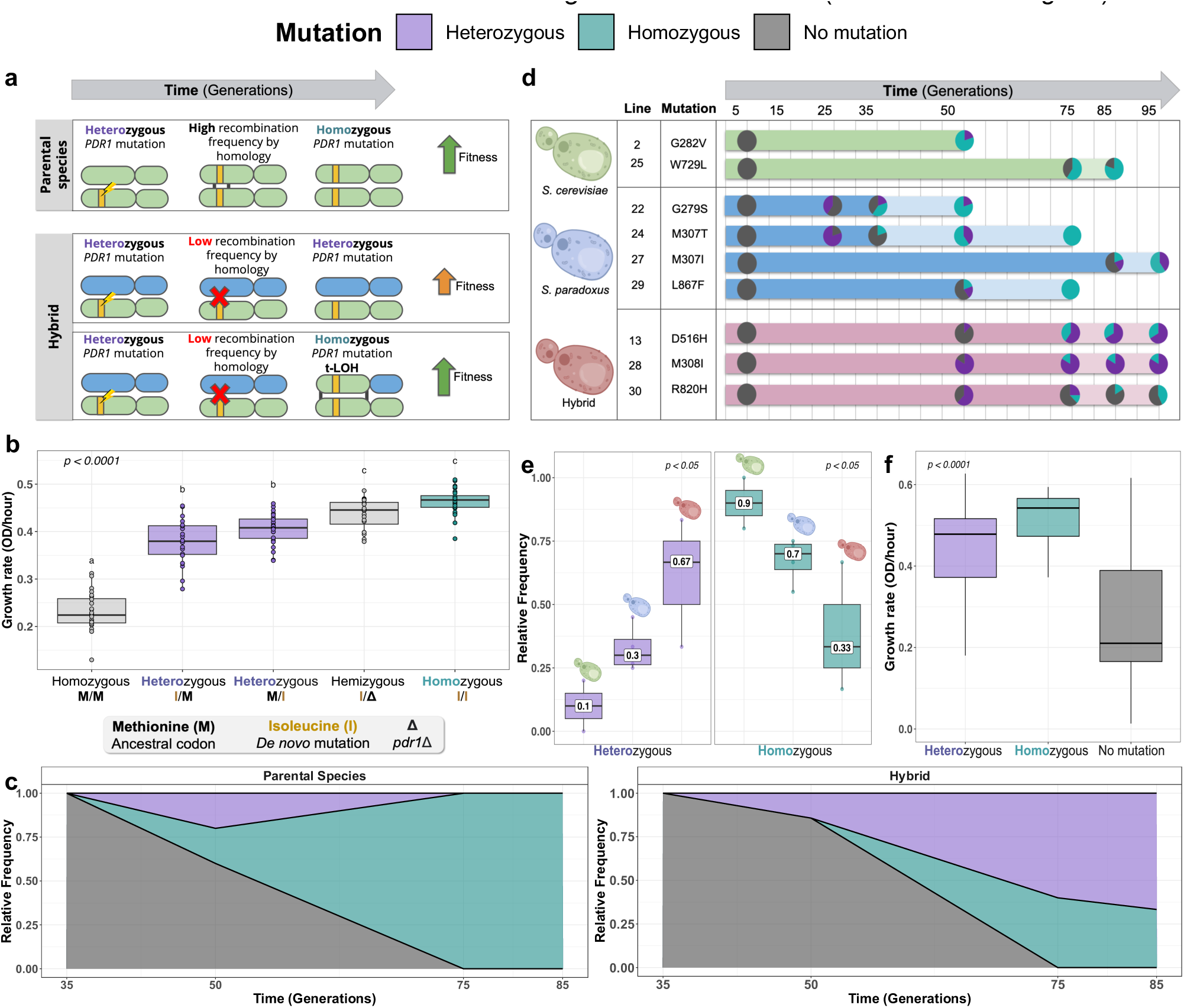
*PDR1* mutations show additive phenotypes such that allele copy number correlates with fitness gain. **a,** Schematic of the hypotheses tested. Rapid homozygosity through LOH allows fitness to increase rapidly in parents. Hybrids achieve homozygosity more slowly and it occurs through major genomic changes in copy number (t-LOH). A single-parent species is depicted for simplicity of visualization. **b,** Growth rate in UV mimetic conditions (4 µM of 4-NQO) of the four types of zygosity (homozygous WT, heterozygous M308I, hemizygous M308I, and homozygous M308I) of *PDR1* constructed by CRISPR-cas9 genome editing (*n* = 22-28 per zygosity). *p*-value for ANOVA test (above) and Tukey post hoc pairwise groups (above each boxplot) are shown. **c,** Temporal dynamics of the evolutionary history of hybrid line 13 and *S. paradoxus* line 29 evolved in UV mimetic conditions. Relative frequencies through time (measured in generations) of the three detected zygosities are shown (heterozygous *PDR1* mutation in purple, homozygous *PDR1* mutation in turquoise, or no mutation on *PDR1* in grey). The relative frequency is displayed from one time point before the first detection of the *PDR1* mutation to up to two time points after. Panel shows a representative example for simplicity but see *Extended Data* Fig. 9 for a comprehensive analysis of all lines (n= 5-10 isolated clones per time point). **d,** Temporal dynamics of the evolutionary history across the three types of zygosity (*n* = 2 lines for *S. cerevisiae*, *n* = 4 lines for *S. paradoxus,* and *n* = 3 lines for hybrid, each line has *n* = 5-10 isolated clones per time point). The dark-colored interval represents the number of generations in which the first homozygous mutation appears. **e,** Relative frequency of homozygous and heterozygous lines across genotypic backgrounds (*S. cerevisiae*, *S. paradoxus,* and hybrid). The relative frequency was determined by calculating the median value of relative frequencies across time points, starting from the first detection of *PDR1* mutations until reaching a homozygous level exceeding 80% or the frequency detected in the last resequenced time point (95 generations). The median values for each genotypic background are highlighted in squares (*n* = 2 lines for *S. cerevisiae*, *n* = 4 lines for *S. paradoxus,* and *n* = 3 lines for hybrid, each line has *n* = 5-10 isolated clones per time point). *p*-value for ANOVA test (above) is shown. **f,** Growth rate in UV mimetic conditions (4 µM of 4-NQO) of hybrid zygosities (heterozygous *PDR1* mutation, homozygous *PDR1* mutation or no mutation on *PDR1*) through time (from isolated clones of the experimental evolution frozen fossils) (*n* = 40-80 isolated clones / line from three hybrid lines evolved in UV mimetic conditions containing *PDR1* mutations: 13, 28 and 30). *p*-value for ANOVA test (above) is shown. Illustrations in **b**, **d**, **e**, and **f** were created with BioRender.com.

Taken together, our findings demonstrate an association between genomic changes leading to the amplification of mutated *PDR1* copies and fitness gains in UV mimetic conditions. Hybrids with *PDR1* mutations do show fitness gains but to a smaller extent than both parental lines. This advantage could derive from the fact that parental genomes achieve higher levels of homozygosity for these mutations (*Fig. 3c*): The LOH rate in hybrids containing *PDR1* mutations was 37.5% (3/8 hybrid lines had a homozygous *PDR1* mutation), whereas it was 100% across the parental *S. cerevisiae* lines (3/3) and 62.5% across *S. paradoxus* lines (5/8), making *PDR1* mutations more visible to selection in parental lines. Thus, the limiting factor may not be the rate and type of mutation but their limited allelic amplification in hybrids (Fig. 4a).

### The Challenge of Attaining Homozygosity Accounts for the Reduced Adaptive Potential of Hybrids

In order to test if Haldane’s sieve slows down adaptation in hybrids, we rely on the following key assumptions: 1) The initial *PDR1* mutation is recessive or incomplete dominant, requiring homozygosity to unlock its full fitness benefits. We thus predict that homozygous *PDR1* mutants will display higher fitness than heterozygous *PDR1* mutants; 2) LOH occurs at a slower pace in hybrid genomes^42,45,91^.

We first validated the adaptiveness of specific *PDR1* mutations. Site-directed mutagenesis was used to introduce seven candidate mutations (G280R, G280S, M308I, and G1042W for *S. cerevisiae*; G279R, G279S, and G281V for *S. paradoxus*) on a plasmid carrying either *S. cerevisiae or S. paradoxus PDR1* gene. After introducing these plasmids into a *S. cerevisiae* strain (BY4741) deleted for *PDR1* (*pdr1*Δ); we found that the mutations conferred significantly higher growth rates in the presence of the UV mimetic drug compared to the wild-type (WT) (*Extended Data* Fig. 8a). These mutations also conferred fitness benefits in the parental backgrounds with slight variations in the extent of effects (*Supplementary Fig. 6*). To further confirm the effects derived from the transcription factor activity of *PDR1*, we measured expression of the downstream drug efflux pump Pdr5^72,73^. We fused Pdr5 to a Green Fluorescent Protein (mEGFP) and measured its expression in the same strain (BY4741) containing *PDR1* mutations on a plasmid. The Pdr5-mEGFP strain exhibited higher fluorescence levels when carrying a plasmid containing specific *PDR1* mutations, compared to wild-type (WT) (*Extended Data* Fig. 8b). Consistent with this observation, we found that the same *PDR1* mutations also lead to resistance to several drugs other than 4-NQO (*Extended Data* Fig. 7c).

To assess the dominance of *PDR1* mutations in a neutral genomic background, independent from potential interference by other mutations as might occur in our lines, we generated homozygous and heterozygous diploid mutants by mating BY4741 and BY4742 haploids and using CRISPR-Cas9 genome editing. We focused on the M308I mutation because it showed the most significant fitness gain and activation of the Pdr5 efflux pump (*Extended Data* Fig. 8). We found that in the homozygous state, this mutation conferred a higher fitness advantage than in the heterozygous state (*Fig. 4b*), confirming its incomplete dominance. The hemizygote also showed a higher growth rate than the heterozygote, suggesting that the mutated allele confers higher benefits in the absence of the WT allele. Taken together, our findings confirm the first key assumption: 1) The initial *PDR1* mutation exhibits incomplete dominance and requires homozygosity to fully contribute to adaptation.

We archived populations regularly over the course of experimental evolution^49^, so we could revive the frozen fossils and isolate some clones to determine the timing of appearance of the various *PDR1* mutations. We revived ∼350 clones from nine hybrid and parental populations that harbor homozygous *PDR1* mutations based on WGS data and sequenced the *PDR1* locus using amplicon Sanger sequencing. We identified three types of zygosity at the *PDR1* locus at intermediate time points of experimental evolution (*Fig. 4c*): heterozygous, homozygous WT, and homozygous mutant. As expected for *de novo* mutations in diploids, mutations were consistently first detected as heterozygous (*Fig. 4d*) but in some cases, the LOH was so rapid that we also detected homozygous mutants (*Fig. 4c left*).

This trend aligns with the increase in fitness recorded during the experimental evolution in their populations of origin but also when analyzing specific isolated clones^49^ (*Extended Data* Fig. 9 and *Extended Data* Fig. 10, respectively). Mutations only became homozygous later (*Fig. 4d*) but with the important difference that the parents become homozygous at a higher frequency than the hybrids. This pattern persisted across all experimental populations: Homozygous genotypes appeared quickly after the initial *PDR1* mutations occurred, and spread rapidly in the parental populations, whereas in the hybrids, even by generation 95, a high proportion of heterozygotes were still observed (*Fig. 4d, see Extended Data* Fig. 9 for detailed analysis). Supporting this trend, we saw in some of the parents (population 2 and 25 of *S. cerevisiae*, population 27 and 29 of *S. paradoxus*) that the emergence of homozygous and heterozygous coincide, indicating that the mechanism of adaptation through LOH can operate quickly (*Fig. 4c* and *4d*). The relative frequency of each mutation in each population through time further shows that the proportion of homozygous mutants was much lower in hybrids compared to parental species (*Fig. 4e*). Remarkably, these proportions (*Fig. 4e*) closely mirror those calculated above from the genome sequences, averaged for each genotypic background (100% vs. 90% in *S. cerevisiae*; 62.5% vs. 70% in *S. paradoxus* and 37.5% vs. 33% in hybrids). To verify that the low homozygote frequency in hybrids was not due to reduced homozygote fitness in the specific hybrid genotypic background, we compared hybrid fitness across generations and populations and confirmed that homozygotes were fitter than heterozygotes and homozygotes WT (*Fig. 4f and Extended Data* Fig. 10 for fitness across generations). These findings validate our final assumption that 2) LOH occurs at a slower pace in hybrids.

## Discussion

Hybridization is a recurring phenomenon in nature that has captured the interest of scientists for decades^1,9,14,92–95^ in fundamental fields but also in applied research such as in agriculture and medical microbiology^24,25,28,96,97^. The adaptive and non-adaptive roles of hybridization have been studied extensively^8,98–107^. However, the negative consequences have mostly focused on reproductive isolation^108,109^ and less on the potential reduction in adaptive rates of hybrids. We previously evolved hybrids of *S. cerevisiae* and *S. paradoxus* species during 100 generations in a stress that mimics UV radiation and observed a reduced adaptive potential of hybrids. We used this system to investigate what could reduce the hybrid rate of adaptation.

Our results reveal that hybrids and parental species have access to the same adaptive changes in key genes. In principle, adaptation can therefore occur through the same mechanisms and at the same rate. We examined the cases of mutations that impacted a transcriptional factor involved in drug resistance because they displayed strong parallelism. The mutations displayed incomplete dominance and could only avoid Haldane’s sieve - the bias against the establishment of recessive beneficial mutations^87–89^ - by achieving homozygosity through LOH. Because LOH depends on recombination and recombination depends on high sequence identity^42,45,91^, this second event (LOH) occurs at a slower rate in hybrids, ultimately contributing to slowing down hybrid adaptation. This is opposite to *de novo* mutations, which accumulate in yeast hybrid genotypes at rates that are not greater than those observed in these parental species^110^. Experiments involving the evolution of heterozygous yeast populations have also shown that LOH frequently unmasks beneficial recessive alleles which can confer significant fitness advantages^90,111–113^. Although not explored here, LOH in hybrids could also limit adaptation in other ways. Since mitotic recombination often extends along the entire length of a chromosome arm^62,64^, especially in heterozygous genomes^63^, an LOH that renders a beneficial mutation homozygous could also bring along other molecular changes or combination of changes that would negatively impact fitness. These could include for instance recessive genetic interactions between the two species that would be revealed following a long LOH^67^.

Our findings contribute to the understanding of the genomic factors shaping asexual microbes. Such hybrids often evolve during domestication, for instance, many beer yeasts are among-species hybrid^114,115^ and these hybrids are known to be largely sterile, i.e. to not have access to sexual reproduction^116^. Even fungal pathogens evolve through recurrent hybridization events^28,97,117–121^ and acquire antifungal resistance and adapt to new hosts with *de novo* mutations and LOH^122–126^. It has indeed been shown that to confer full resistance to antifungals, a mutation in a transcriptional regulator needed to be followed by LOH^127^. Antimicrobial resistance has also been shown to depend on an LOH event in *S. cerevisiae,* in order to render a loss-of-function mutation homozygous^128^. Understanding which conditions could slow down the rate of LOH, such as heterozygosity along the chromosome as we exemplify here, or the linkage to other potentially deleterious mutations^129^, is there key to understanding evolution in an applied context such as antimicrobial resistance. Other asexual cells that evolve in a similar manner are somatic cells. Cancer cells reproduce somatically and usually evolve by LOH, since most of the mutations associated with tumor progression need to remove the dominant alleles of tumor suppressors to become active^130,131^. The phenomenon we uncovered here, whereby some genotypes experience lower rates of LOH, thus has also consequences that extend beyond the study of hybrids.

## Methods

### Experimental Crosses and Previous Experimental Evolution

The *S. cerevisiae* and *S. paradoxus* strains used were described in^49^ and were derived from the natural strains LL13_054 and MSH-604 isolated in North American forests^13,48^ (*Supplementary Table S2*). To prevent mating type switching in haploids, the *HO* locus was replaced with resistance cassettes (HPHNT1 for Hygromycin B resistance and NATMX4 for Nourseothricin resistance)^132^ through homologous recombination as described in *Table S1*. A total of 90 experimental strains were constructed by crossing haploid strains (30 *S. cerevisiae*, 30 *S. paradoxus* and 30 hybrid), so that each starting parental and hybrid diploid population is the result of an independent mating event as described in ^49^. The 90 populations were evolved for 100 generations (*Fig. 1a*) as described in^49^. Briefly, we used a non-DNA damaging growth condition called control (YPD 1% yeast extract, Fisher BioReagents™, USA; 2% tryptone, BioShop^®^, Canada; and 2% D-glucose, BioShop^®^, Canada) and a DNA damaging growth condition supplemented with a UV mimetic molecule^50^ 4-Nitroquinoline 1-oxide (4-NQO) (Sigma–Aldrich, cat. no. N8141, batch #WXBC3635V, Canada)^50^.

### Fitness Assays on Individual Clones

Growth assays were conducted on individual clones, which were used for genome sequencing and isolated from the glycerol stocks from experimental evolution^49^ (*Supplementary Table S1*). Ancestor strains (*n* = 90) as well as the populations evolved in YPD (*n* = 90) and in YPD + 4-NQO (*n* = 90) were pre-cultured in 1 mL of YPD in 96 deep-well plates and incubated for 24 h at 25 °C. Subsequently, 20 µL of these pre-cultures were grown in 96-well flat-bottomed culture plates in 180 µL of medium (YPD or YPD + 4 µM of 4-NQO), resulting in an initial OD_595_ of approximately 0.1. A transfer cycle was performed at 24 h after approximately ∼ 5 generations in rich conditions. Each culture was diluted approximately 30-fold by transferring 6 µL of grown culture into 194 µL of fresh medium to initiate a new round of growth at an OD_595_ starting at about 0.03. Incubation at 25 °C was performed directly in three temperature-controlled spectrophotometers (Infinite^®^ 200 PRO, Tecan, Reading, UK) that read the OD_595_ at intervals of 15 min throughout the two cycles performed. All samples were randomized across plates, temperature-controlled spectrophotometers and days.

### DNA Extraction, Library Construction and Whole Genome Sequencing

We obtained whole-genome sequences of 270 individual clones derived from the 270 experimental lines^49^ (*Supplementary Table S1*). We extracted genomic DNA from overnight YPD cultures derived from each clone according to the manufacturer’s instructions (MasterPure™ Yeast DNA Purification Kit, Biosearch Technologies - Lucigen, Wisconsin, USA) and purified on Axygen™ AxyPrep Magnetic PCR Clean-up SPRI beads (Axygen Inc, New York ,USA). Five DNA libraries were prepared using RIPTIDE™ High Throughput rapid DNA library prep in 96-well plate format (iGenomX, South San Francisco, USA)^133^. The quality of the libraries was verified using an Agilent BioAnalyzer 2100 electrophoresis system (Genomic Analysis Platform of the Institute of Integrative Biology and Systems of Université Laval, Quebec, Canada). Pooled libraries were sequenced using paired-end 150 bp reads on different lanes of an Illumina NovaSeq 6000 (Illumina, San Diego, USA) at the Genome Quebec Innovation Center (Montreal, Canada).

### Flow Cytometry Analysis of Ploidy

DNA content was measured by flow cytometry using the SYTOX™ green staining assay (Thermo Fisher, Waltham, USA) as in ^42,48^. Haploid and diploid strains of the *S. cerevisiae* isolate LL13_054 were used as haploid and diploid controls, respectively. As triploid control, we used a cross between *S. paradoxus* subspecies B (MSH-604) and *S. paradoxus* subspecies C (LL11_004) strains. As tetraploid control, we used a cross between *S. paradoxus* subspecies B (91_202) and *Saccharomyces cerevisiae* (LL13_054) strains^48^. Because we do not have controls from each genetic background, there may be slight differences in DNA content measurements and thus we inferred ploidy between our lines and the controls. The 270 individual clones derived from the 270 experimental lines (*Supplementary Table S1*) from ^49^ and used for whole-genome sequencing were thawed from glycerol stocks and grown on solid YPD omnitray plates (25°C, 72 h). They were inoculated in 1 mL of YPD in 96 deepwell plates and incubated for 24 h at 25 °C. Cells were subsequently prepared for flow cytometry as in ^134^. They were fixed in 70% ethanol and kept frozen at - 20 °C for further analysis. RNA was eliminated using 0.25 mg mL−1 RNase A during an overnight incubation at 37 °C. Cells were washed twice with sodium citrate (50 mM, pH 7) and stained with a SYTOX™ green concentration of 0.6 μM for 1 h at 25 °C in the dark. Cell concentration was adjusted in sodium citrate (50 mM, pH 7) to be less than 500 cells/μL. Five thousand cells from each of the 300 samples were analyzed in 96-well plates in a CytoFLEX Platform flow cytometer (Beckman Coulter, California, USA) at the Feldan Therapeutics facility (Quebec, Canada). Cells were excited with the blue laser at 488 nm and fluorescence was measured in a green fluorescence detection channel (525/40 nm). The distributions of the green fluorescence values were processed to find the two main density peaks, which correspond to the two cell populations in G1 and G2 phases, respectively. DNA content value was calculated as a median of the fluorescence of the two main density peaks.

### Quality Assessment and Read Mapping of Next-Generation Sequencing Data

Raw reads from barcoded samples of the five libraries were demultiplexed using DemuxFastqs from fgbio tools^135^ v1.5.0 (*Supplementary Table S3*). Reads were trimmed using Trimmomatic^136^ v0.36 with parameters ILLUMINACLIP:Trimm_seqs.fa:6:20:10 and using Trimm_seqs.fa (*Source Data 1*) as a list of adapter sequences used. To assess the quality of both pre- and post-trimming sequencing reads, we used FastQC v0.11.9 ^137^ and MultiQC v1.11^138^.

Reads from *S. cerevisiae* samples were mapped on the indexed reference genome of *S. cerevisiae* strain YPS128^139^, which in our study is named LL13_054, and *S. paradoxus* samples were mapped on the *S. paradoxus SpB* (named MSH-604) genome^139,140^. Reads from hybrid lines were mapped on a concatenated genome comprising the two respective parental genomes end to end. The BWA-MEM algorithm^141^ v0.7.17 was used for mapping. Mapped reads were processed by genome-sorting algorithms using samtools v1.8^142^ and quality was assessed by mapping coverage with goleft v0.2.2^143^. The average mean read depth across samples was about 100X (*Supplementary Fig. 1*). We excluded line 16 from the *S. cerevisiae* population evolved under UV mimetic conditions due to its low quality. We used Picard tools v2.26.11^144^ for adding Read Groups groups with AddOrReplaceReadGroups, and we removed duplicate reads with MarkDuplicates with parameter REMOVE_DUPLICATES = true.

### Analysis of Read Depth (Aneuploidies and Loss of Heterozygosity)

Mean read depth over 1 kbp windows were obtained with BamStats04 from Jvarkit tools v2021.08.10^145^ and makewindows from bedtools tools v2.30.0^146^. We first eliminated some sequences of the hybrid lines evolved in UV mimetic because the sequencing was weak (lines 1 and 21) or because the content of one of the parental genomes was the majority (lines 10 and 25) (*Supplementary Fig. 7, 8 and 9*). We also verified the results of **Flow Cytometry Analysis of Ploidy** section and compared DNA content by measuring average read depth across genomes (*Supplementary Fig. 10*). We computed the median chromosome read depth and the median whole genome read depth for each line. In order to detect the number of gained or lost chromosomes, we divided each chromosome’s median read depth by the genome-wide median read depth. We standardized this value by the value of the corresponding ancestors to obtain the relative read depth (log2 fold change). Values with a chromosome median read depth higher than the genome-wide median represent gains in DNA content (gradient towards red in *Fig. 2a*, *Fig. 2c* and *Extended Data* Fig. 3) and values with a median read depth lower than the genome-wide median represent losses in DNA content (gradient towards purple in *Fig. 2a*, *Fig. 2c* and *Extended Data* Fig. 3) for each individual chromosome. We also computed the number of chromosomes with aneuploidies per line by considering an aneuploidy as a deviation (increase or decrease) of 30% with respect to the genome-wide median read depth as in ^26^. The positional coverage mapping of hybrid genomes unveiled terminal regions with pronounced increases in read depth in one parental chromosome copy and concurrent decreases in the other copy (*Extended Data* Fig. 3), revealing the presence of reciprocal crossovers between chromosomes. These tracts that extend to the telomeres and usually measure between 50-100 kb correspond to Terminal-Loss of Heterozygosity (t-LOH) regions^62,63,66^. To quantitatively assess the number of t-LOH events in hybrid lines, we identified regions with simultaneous increases and decreases in read depth (deviation of 30% with respect to the genome-wide median read depth) in both chromosomal copies exceeding a size threshold of 20 kb.

### Functional Analysis of *de novo* Mutations

SNP calling was performed with Haplotype Caller (gatk-v4.1.4.1)^147,148^. Before generating the GVCFs, we added a RG (read group) tag to individual BAM files. After SNP calling, genotyping of GCVFs was performed with GenotypeGVCFs. For variant filtration, we applied standard hard filters with options: QUAL by depth (QD) < 2.0, mapping quality (MQ) < 40.0, Fisher’s exact tests of strand bias (FS) > 60.0, symmetric odds ratio test of strand bias (SOR) > 3.0, mapping quality rank sum test (MQRankSum) < -12.5, rank sum test for site position within reads (ReadPosRankSum) < -8.0, Genotype Quality (GQ) < 20, and Coverage (DP) < 3. We selected variants which passed the above filters and we excluded INDELS focusing exclusively on substitutions (SNP) for subsequent analysis. We excluded the pre-existing genetic variation with respect to the reference genome, by removing any variant that was already present in the ancestral strain for each evolved line (*Extended Data* Fig. 5). For annotation purposes, we used the *S. cerevisiae* genome assembly R64-1-1 (*Saccharomyces cerevisiae* S288c assembly from *Saccharomyces* Genome Database, INSDC Assembly GCA_000146045.2, Sep 2011) for *S. cerevisiae* genomes. Subsequently, we generated maps annotations with Liftoff (v1.6.3)^149^ of *S. paradoxus* genome from the *S. cerevisiae* genome assembly R64-1-1 (*Saccharomyces cerevisiae* S288c assembly from *Saccharomyces* Genome Database, INSDC Assembly GCA_000146045.2, Sep 2011). We finally generated a combined annotated genome for hybrid analysis. The variants were ultimately annotated using Ensembl Variant Effect Predictor (VEP) v110^150^. We examined missense variants to perform Gene Ontology (GO) using bioMart^151^ and clusterProfiler^152^.

### Validation of the Adaptiveness of Mutations

We used ChimeraX v1.5^153^ to visualize Pdr1p amino acid changes we found throughout the experimental evolution and other mutations found in the literature^75–79^ (*Source Data 2*) on the AlphaFold^154,155^ generated structure for Pdr1p (AF-P12383-F1, *Source Data 3*). Subsequently, we analyzed Pdr1 *Nakaseomyces glabratus* protein by superimposing structures and amino acid changes^80,83,84^ across species (*Source Data 4;* AF-B9VI40-F1, *Source Data 5*). We next used a set of plasmids derived from MoBy-ORF library, in which genes are controlled by its native promoter and terminator^156^, to express the *PDR1* sequences from either *S. cerevisiae* (BY4741) or *S. paradoxus* (MSH-604). Candidate mutations (G280R, G280S, M308I, and G1042W, being G1041W in LL1304 strain for *S. cerevisiae* sequence, G279R, G279S, and G281V for *S. paradoxus* sequence) were inserted by site-directed mutagenesis. As controls, we used the plasmid without the *PDR1* gene cloned (Empty) or the plasmid containing the Wild-Type (WT) *PDR1* sequence. We introduced these plasmids following a modified lithium acetate transformation protocol ^157^ in a *S. cerevisiae* lab strain BY4741, and natural strains *S. cerevisiae* LL13_054 and *S. paradoxus* MSH-604 in both WT and *pdr1*Δ (previously constructed by replacing *PDR1* locus with a NATMX4 module) backgrounds. Fitness assays were performed following the same steps as described above in the Fitness **Assays on Individual Clones** section. We added extra conditions to the ones previously used (4 µM of 4-NQO): 8 µM and 10 µM of 4-NQO (*Supplementary Fig. 6*). We also conducted an assay on the *S. cerevisiae* lab strain BY4741 *pdr1*Δ containing the same mutations on the same plasmids and exposed them to antifungal azoles. After adjusting cell density to an OD_595_ of 1, we made three serial dilutions ⅕ in 200 μL of water (40 μL of cells in 160 μL of water). We spotted 5 μL of each dilution on YPD + 0.2% DMSO, as a control, YPD + 16 μg/mL Fluconazole (FLC), YPD + 2 μg/mL Itraconazole (ITR) or YPD + 0.5 μg/mL Voriconazole (VRC) and incubated at 30 °C for 48h. We assessed the expression of the downstream Pdr5p drug efflux pump by fusing Pdr5 to a Green Fluorescent Protein (mEGFP) (Pdr5-mEGFP) in a BY4741 *pdr1Δ* lab strain expressing *PDR1* mutants from the above described pMoBY plasmids. For a comprehensive list of strains refer to *Supplementary Table S2* and for a comprehensive list of oligonucleotide sequences refer to *Supplementary Table S4*.

### Incomplete-Dominance Assay

To evaluate the dominance of the *PDR1* mutations, we created *S. cerevisiae* diploids harboring either homozygous or heterozygous M308I substitutions using CRISPR-Cas9 genome editing and mating strategy involving BY4741 and BY4742 haploids (*Supplementary Table S2*). We replaced the *PDR1* locus with NATMX4 (in BY4741) or HPHNT1 (in BY4742) modules specifically targeted by two different gRNA using a modified protocol from ^158^. Yeast cells were transformed following a modified lithium acetate transformation protocol ^157^ with a pCAS-NAT or pCAS-HPH plasmid (Addgene plasmid 6084747 modified by ^159^ and ^160^ using the same approach as in ^161^) expressing both the gRNA of interest (NATMX4 or HPHNT1), the *Streptococcus pyogenes* Cas9 ^162^ and a donor DNA sequence featuring 40 bp homology arms flanking the *PDR*1 DNA sequence. The donor DNA sequences were WT (*PDR1*), the mutation desired (M308I) or a stop codon in the first methionine (M1Stop) (Oligonucleotide sequences can be found in *Supplementary Table S4*). Finally, we mated BY4741 and BY4742 haploids with the desired mutations to create diploids WT homozygous (*pdr1Δ*::*PDR1/pdr1Δ*::*PDR1*), M308I homozygous (*pdr1Δ*::*PDR1*(M308I)*/pdr1Δ*::*PDR1*(M308I)), heterozygous (*pdr1Δ*::*PDR1/pdr1Δ*::*PDR1*(M308I) *or pdr1Δ*::*PDR1*(M308I)/*pdr1Δ*::*PDR1*) or hemizygous (*pdr1Δ*::*PDR1*(M308I)/*pdr1Δ*::M1Stop) (refer to *Supplementary Table S2* for a comprehensive list of strains).

### Allele frequency dynamics on *PDR1* mutants

Whole population samples were archived regularly during the evolution experiment^49^. From these, we isolated clones to estimate the timing of the appearance of the various *PDR1* mutations and to track the dynamics between homozygous and heterozygous zygosities in the population. We revived approximately 350 clones from nine lines (*Supplementary Table S1*) and sequenced the *PDR1* locus using amplicon Sanger sequencing (Oligonucleotide sequences can be found in *Supplementary Table S5*). We designed specific primers for each mutation (Oligonucleotide sequences can be found in *Supplementary Table S4*) and used PCR to amplify those regions. We performed an exhaustive zygosity analysis across time points (each time point represents five generations) to quantify the ratio of homozygous to heterozygous variants. We sequenced clones from the first detection of *PDR1* mutations until reaching a homozygous frequency exceeding 80%, or if this does not occur, we sequenced up to time point 19 (95 generations). Fitness assays in hybrid lines were performed following the same steps as described above in **Fitness Assays on Individual Clones**.

### Data availability

Supplementary material including Supplementary tables and Source data can be found at Zenodo^163^ (https://zenodo.org/records/10389558). Sequencing data is accessible at the NCBI Sequence Read Archive (SRA) under BioProject PRJNA1045261. The demultiplexing process details are available at Zenodo^163^ and *Supplementary Table S3*. Tables and scripts for figure generation can be found at Zenodo^163^ and GitHub (https://github.com/cbautistaro/Bautista2023_LOH_project). PDB-formatted files of the AF2-generated models can be found at Zenodo^163^. Strains are available upon request. Data was analyzed using bash and R version 4.2.0.

## Acknowledgments

We thank Nadia Aubin-Horth for feedback on the manuscript, Alexandre Dubé for assistance in the laboratory work, Angel Cisneros, Lorena Ament, Alexandre Andreas Rego, Damien-Biot Pelletier, Adarsh Jay, and David Bradley for help with data analysis and members of the Landry Lab for helpful discussions.

## Author’s contributions

CB, IGA, and CRL designed the research. CB performed the experiments, collected, and analyzed the data with the assistance of IGA, MU, AF, DB, RS and CRL. CB wrote the manuscript with the assistance of CRL. CB, IGA, MU, AF, DB, RS, and CRL edited the manuscript. CRL was responsible for funding acquisition. All authors read and approved the final manuscript.

---

**Extended Data Fig. 1.**
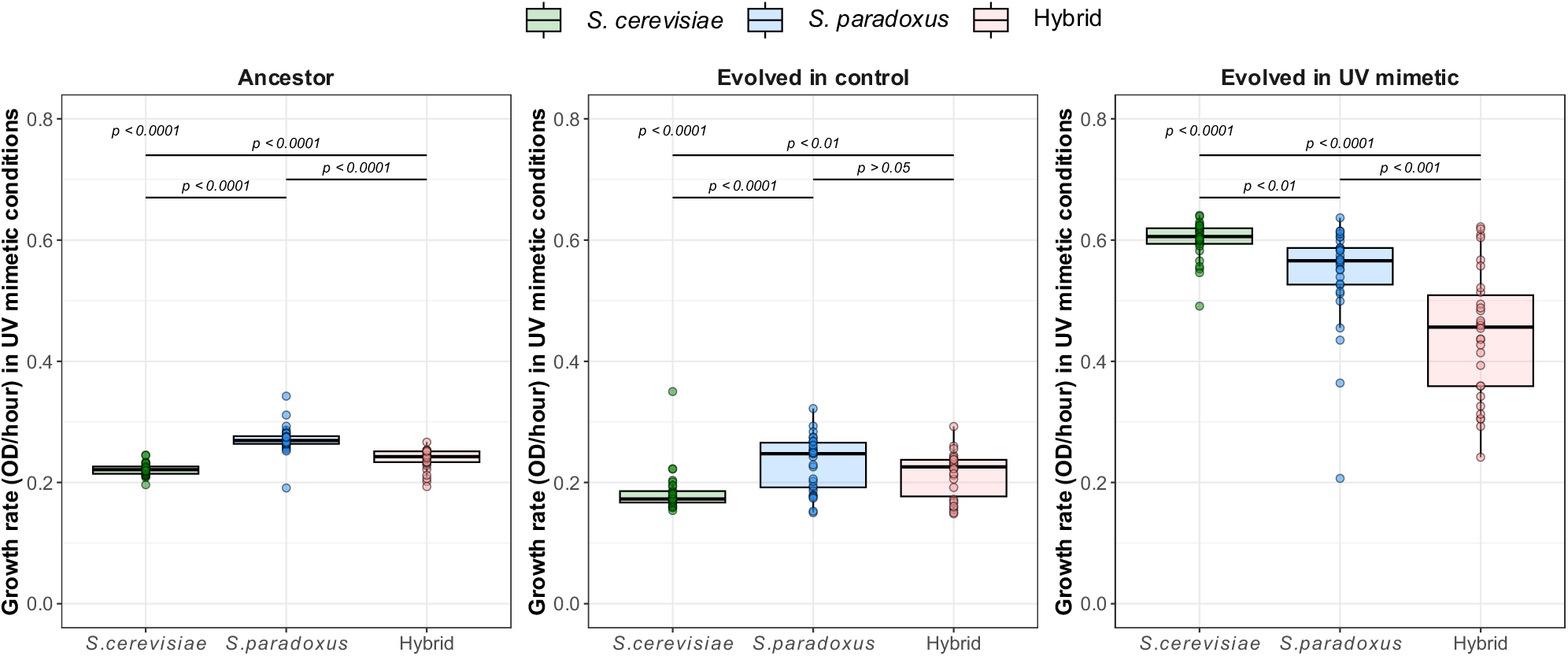
Growth rate of evolved lines in UV mimetic conditions. Growth rate of lines from isolated clones derived from each ancestor population and from each evolved population in control or UV mimetic conditions in the presence of a UV mimetic (4 µM of 4-NQO) (*n* = 30 lines for each genotypic background and condition). *p*-values for ANOVA (above) and t-test for paired lines (below) are shown.

**Extended Data Fig. 2.**
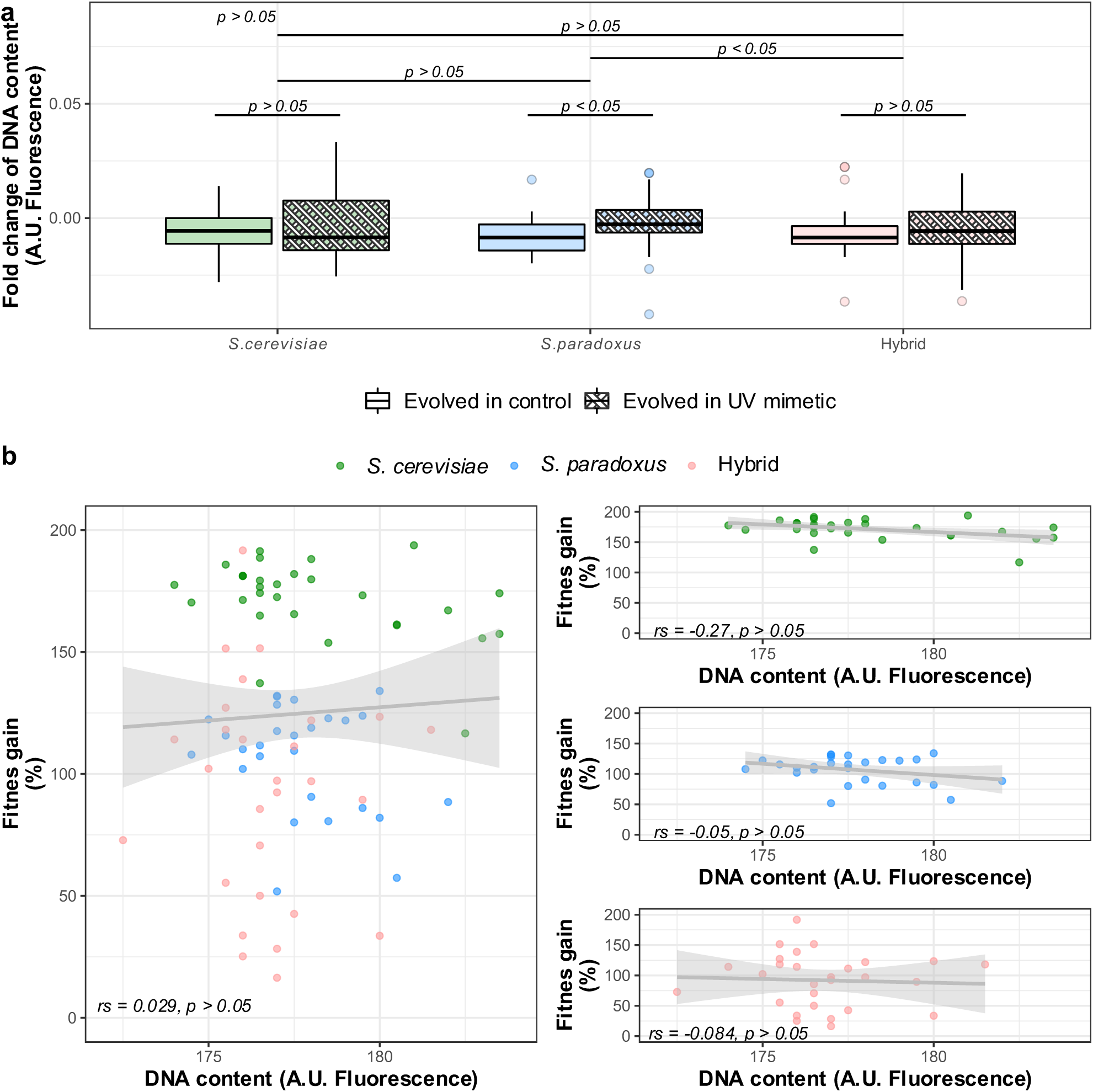
Low frequency of ploidy changes during experimental evolution. **a**, Fold change of DNA content is shown for each genotypic background and condition (*n* = 30 lines for each genotypic background and condition). Fold change of DNA content is represented by the natural log of the ratio between the median DNA content of the lines evolved in UV mimetic conditions and that of the ancestor lines. *p*-values for ANOVA (above) and t-test for paired lines and conditions are shown (*n* = 30 lines for each genotypic background and condition). **b,** Fitness gain (% change in growth rate between initial and final time points) under UV mimetic conditions is not correlated with ploidy level. Ploidy is measured as the median of the distance surrounding the fluorescence peaks (G1 and G2 cell cycle phases) of DNA content (A.U. Fluorescence). The correlation for the three genotypic backgrounds and the individual correlations for each genotypic background are shown. Spearman’s rank coefficients (rs) and associated *p*-values are shown (*n* =30 lines for each genotypic background). A.U. refers to arbitrary units.

**Extended Data Fig. 3.**
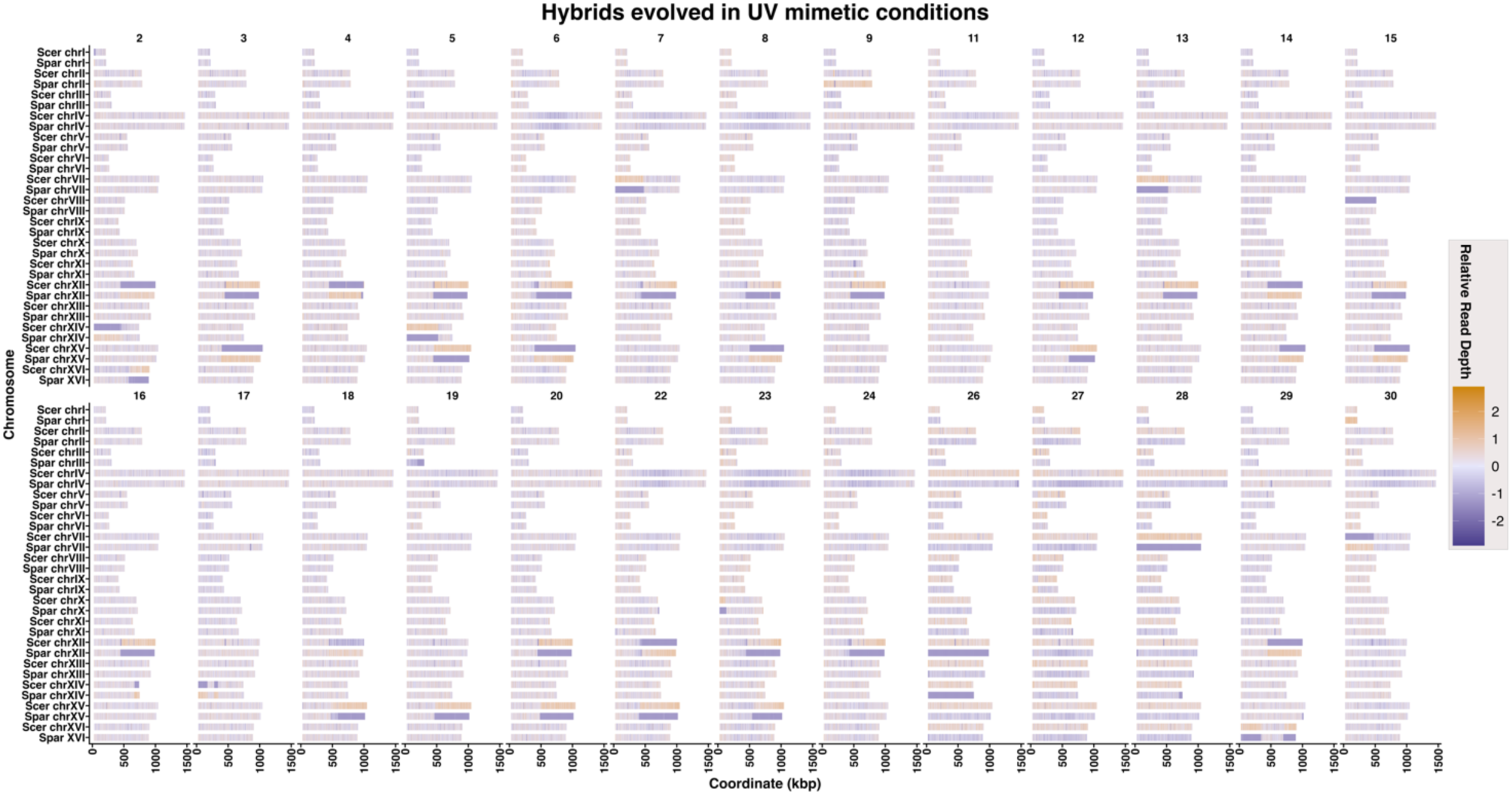
Display of read depth variation across chromosomes for hybrid lines. Detection of Terminal LOH (t-LOH) through simultaneous increases and decreases in read depth (deviations of 30% from the genome-wide median read depth). Lines 26, 27, and 28 are non-diploid genomes showing an increased copy of *S. cerevisiae* genome across all chromosomes.

**Extended Data Fig. 4.**
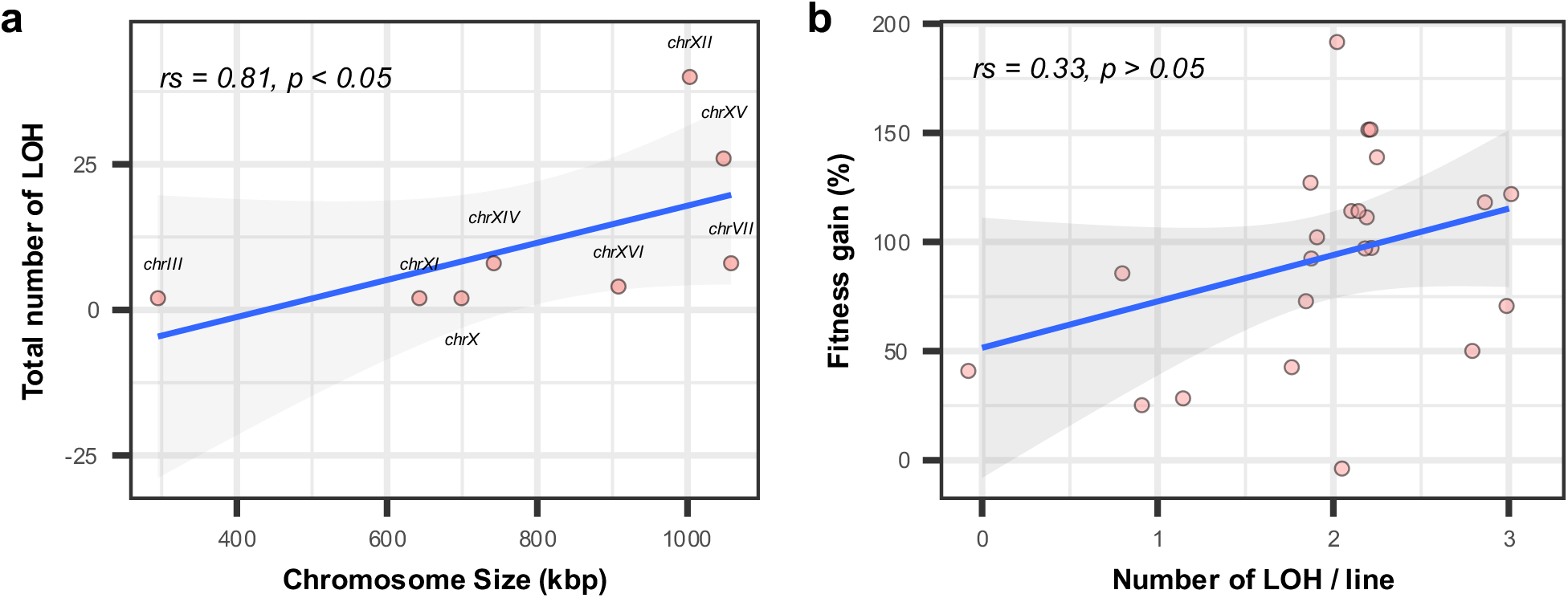
LOH dynamics across hybrid lines in UV mimetic conditions. **a,** Total number of LOH correlates with chromosome size (kbp). Spearman’s rank correlation coefficients (rs) and *p*-values are shown for hybrids evolved in UV mimetic conditions (*n* = 8 chromosomes, the ones affected by t-LOHs). **b,** Fitness gain (% change in growth rate between initial and final time points) as a function of the number of t-LOHs/line. Spearman’s rank correlation coefficient and *p*-value are displayed (*n* = 26 hybrid lines).

**Extended Data Fig. 5.**
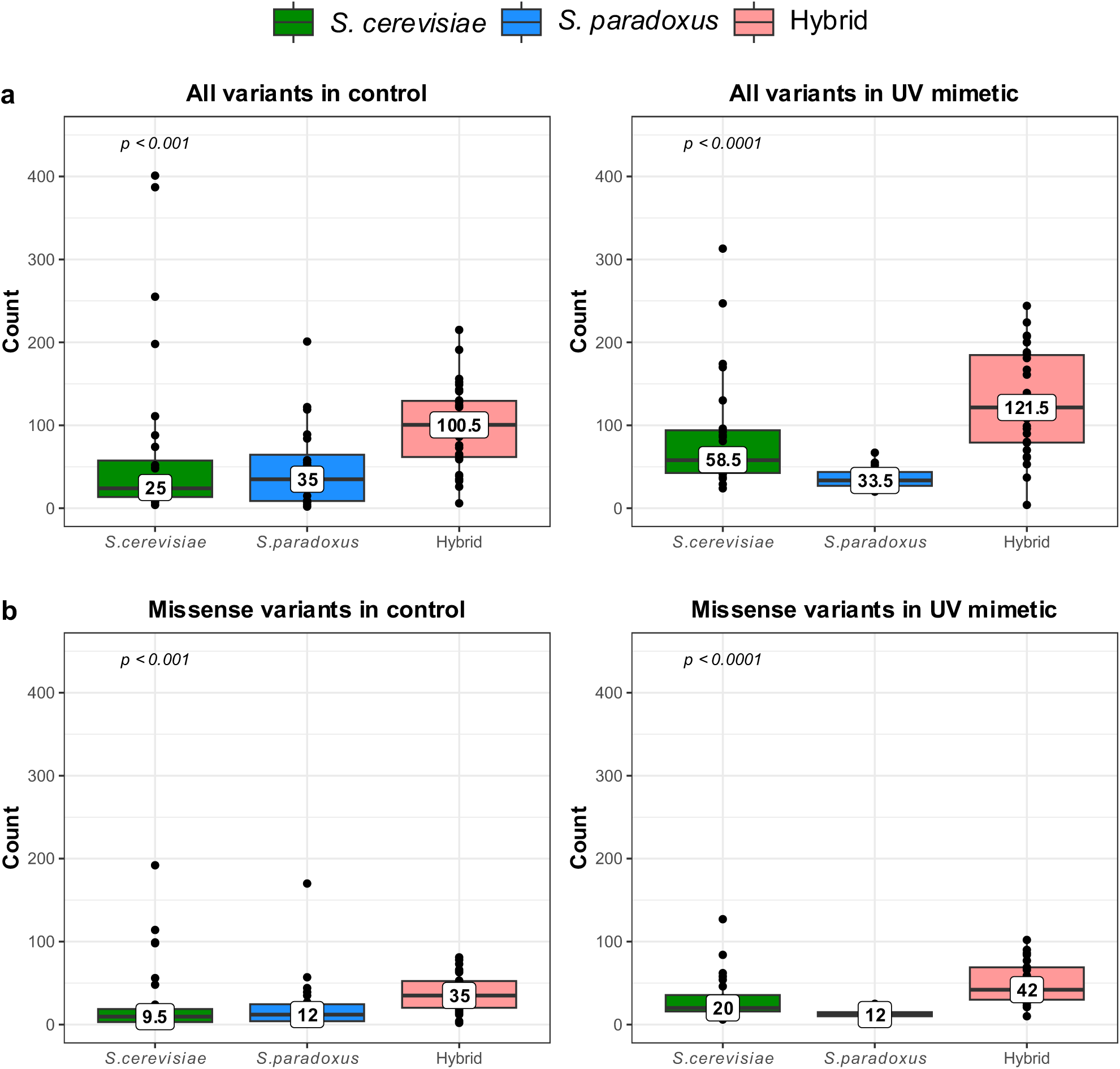
SNPs count through genotypic backgrounds. **a,** Count of SNPs across genotypic backgrounds in both control and UV mimetic conditions. **b,** Count of total missense SNPs across genotypic backgrounds in both control and UV mimetic conditions (n=30 lines per genotypic background). *p*-value for Kruskal-Wallis test (among genotypic backgrounds) is shown.

**Extended Data Fig. 6.**
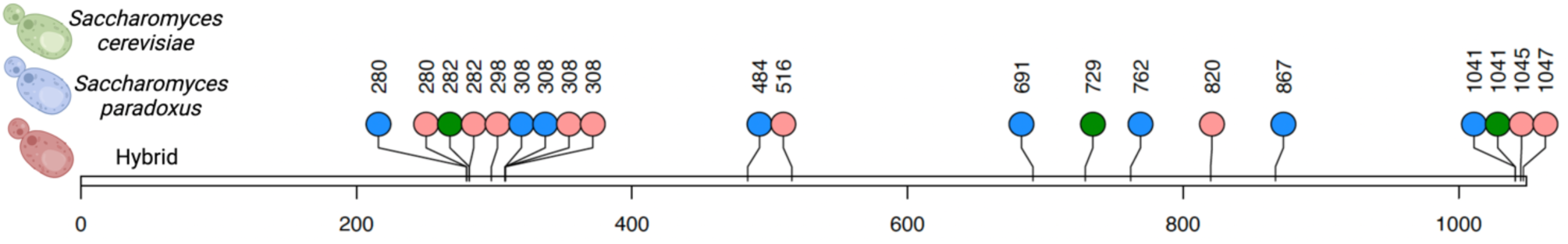
Amino acid changes in Pdr1 protein. Identified non-synonymous mutations are represented along Pdr1p. Each data point is a mutation and numbers represent the amino acid position. Illustration was created with BioRender.com.

**Extended Data Fig. 7.**
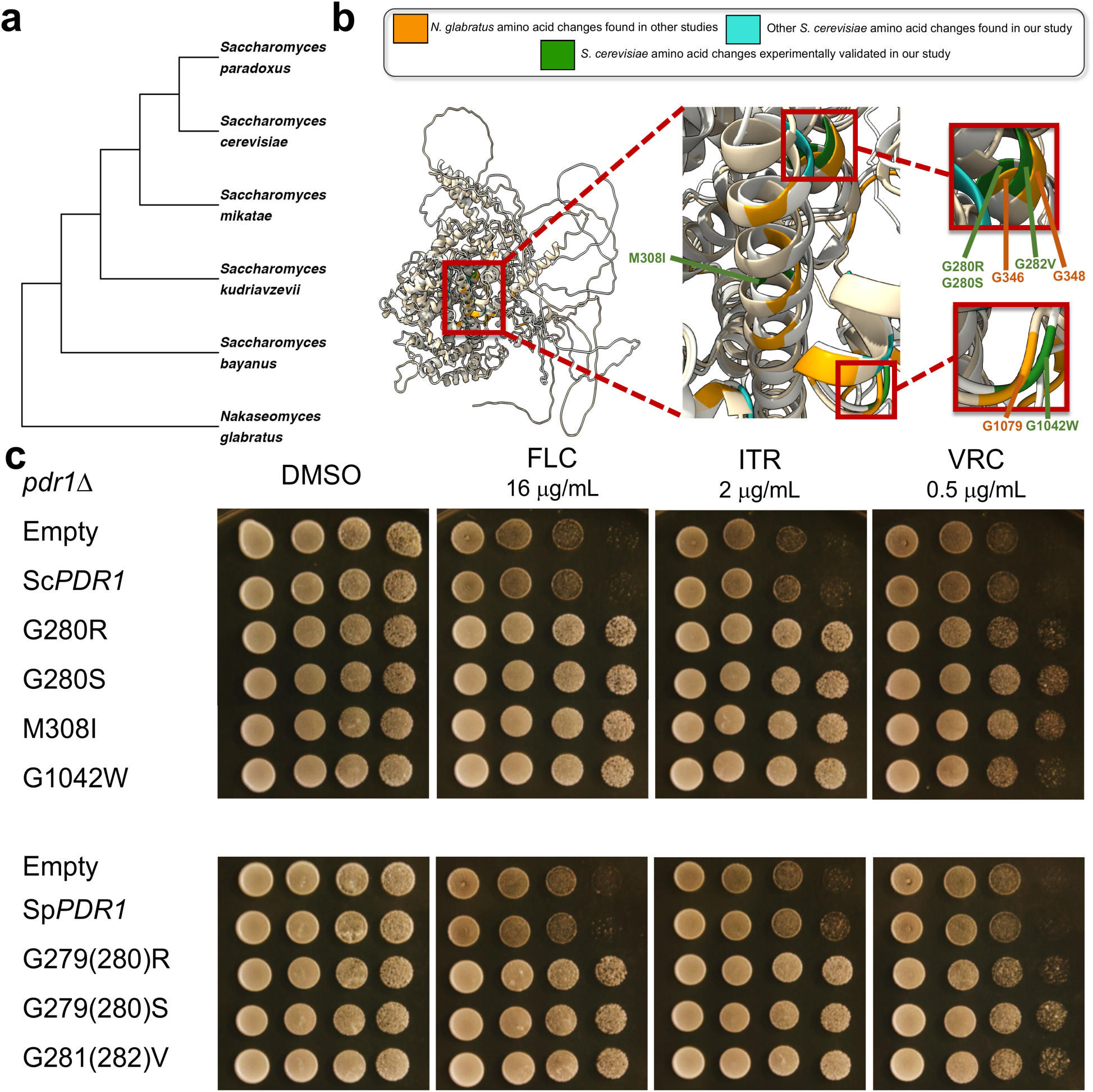
Functional analysis of *PDR1* mutations in *S. cerevisiae* and *Nakaseomyces glabratus*. **a,** Phylogenetic tree showing the relationships between *S. cerevisiae*and *N. glabratus*^164^. **b,** Pdr1p structure aligned between *S. cerevisiae* (white colored) and *N. glabratus* (cream colored) modeled with AlphaFold featuring amino acid changes identified in this study, alongside amino acid changes reported in the literature for *N. glabratus*^80–86^. Cluster of amino acid changes is shown in the insets. **c,** Spot assay of the *S. cerevisiae* BY4741 *pdr1*Δ lab strain expressing *PDR1* variants from a pMoBY plasmid in which the *PDR1* gene is controlled by its native promoter and terminator^156^. This plasmid contains *PDR1* gene from *S. cerevisiae (*Empty: plasmid without the *PDR1* gene; Sc*PDR1*: Wild-Type; G280R; G280S; M308I or G1042W) or from *S. paradoxus* (Empty: plasmid without the *PDR1* gene; Sp*PDR1*: Wild-Type; G279R; G279S or G281V). Growth conditions were DMSO (as a control) and antifungal azoles: Fluconazole (FLC), Itraconazole (ITR), and Voriconazole (VRC).

**Extended Data Fig. 8.**
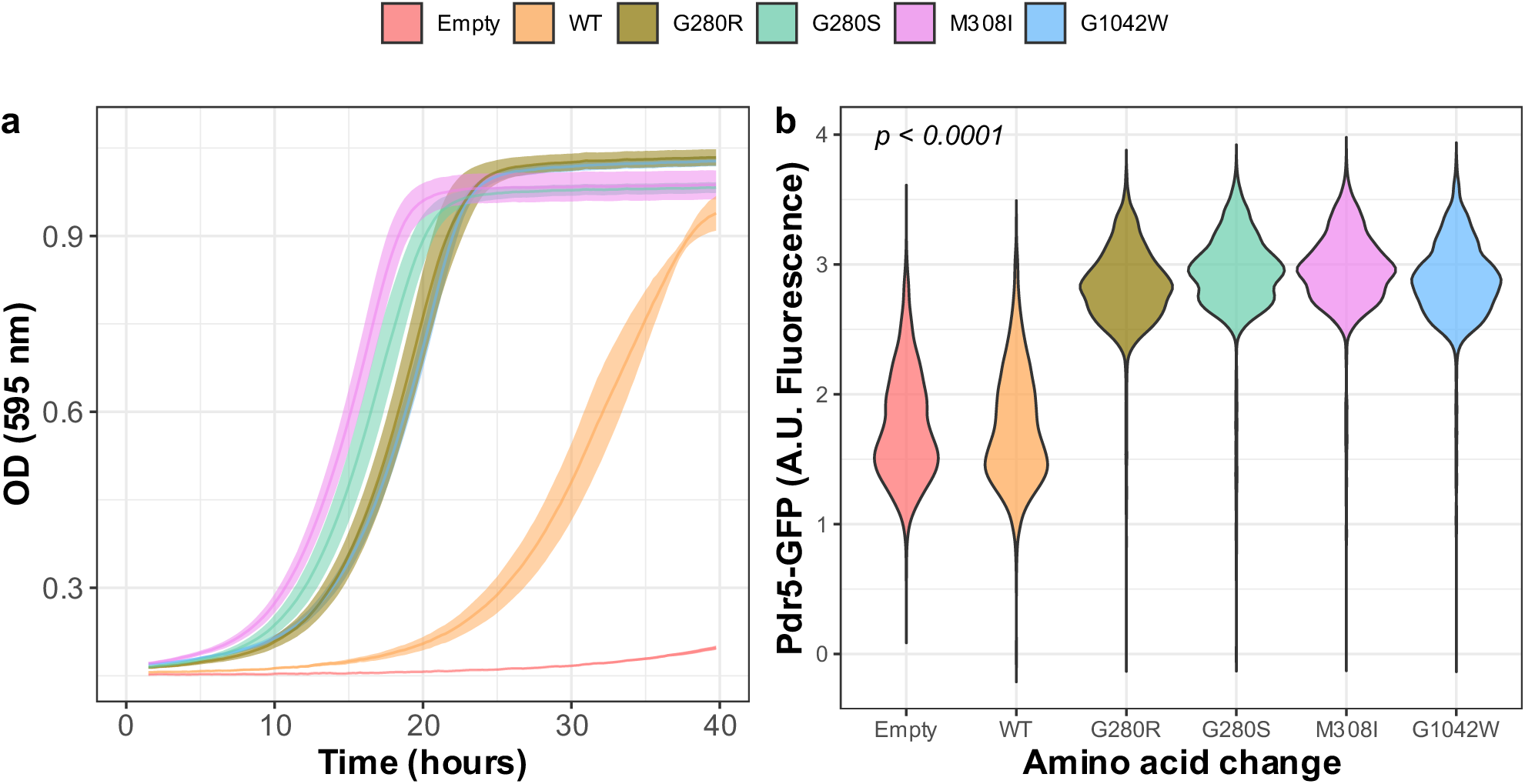
Functional analysis of *PDR1* mutations. **a,** Optical density as a function of time in UV mimetic conditions (4 µM of 4-NQO) of the *S. cerevisiae* BY4741 *pdr1*Δ lab strain expressing *PDR1* variants from a pMoBY plasmid in which the *PDR1* gene is controlled by its native promoter and terminator^156^. **b,** Distribution of cell fluorescence (A.U. Fluorescence) in the population of cells (BY4741 *pdr1*Δ) expressing Pdr5-mEGFP and *PDR1* various mutants from a pMoBY plasmid (*n* = 6). As controls, we used the plasmid without the *PDR1* gene cloned (Empty) or Wild-Type (WT), which is the plasmid containing the *PDR1* gene without mutation. A.U. refers to arbitrary units.

**Extended Data Fig. 9.**
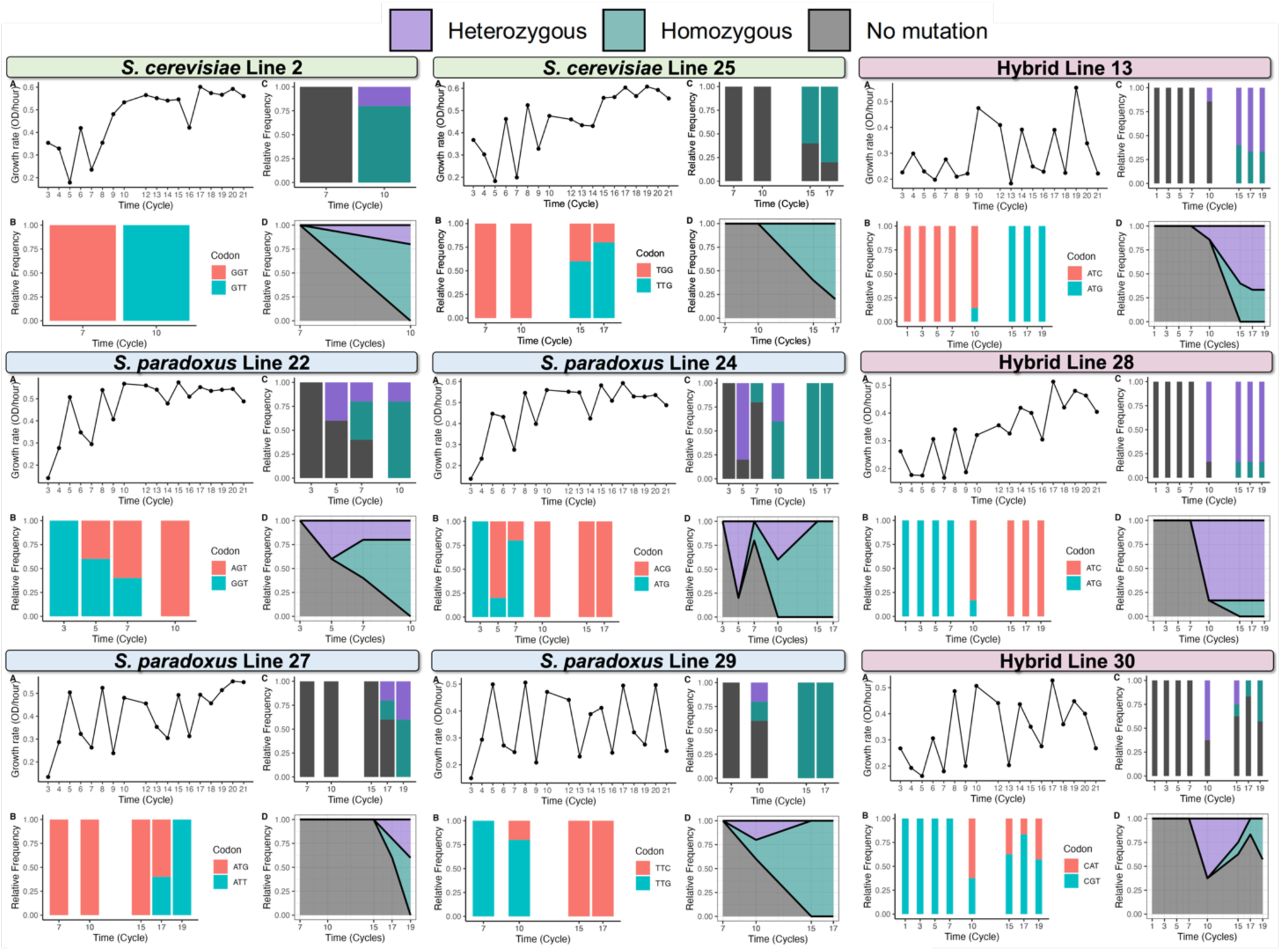
Temporal dynamics of the evolutionary history of *PDR1* mutations across lines. Sanger re-sequencing of individual isolated clones across experimental evolution times points to assessing the *PDR1* genotype. For each line we show: a) Fitness gain through experimental evolution^49^ of our pool of individuals, in which growth rate (OD/hour) is presented across time (time points). b) Relative frequency across time (time points) of ancestor or evolved codon. c) Relative frequency across time (time points) of each different background identified for the *PDR1* locus: heterozygous (*PDR1* mutation present on only one chromosome), homozygous (resulting from an LOH event where both chromosomes have the same *PDR1* mutation), and ancestral WT codon (no *PDR1* mutation detected). d) Expansion across time (time points) of each different background identified for the *PDR1* locus. Each time point represents five generations^49^ (*n* = 2 lines for *S. cerevisiae*, *n* = 4 lines for *S. paradoxus,* and *n* = 3 lines for hybrid, each line has *n* = 5-10 isolated clones per time point).

**Extended Data Fig. 10.**
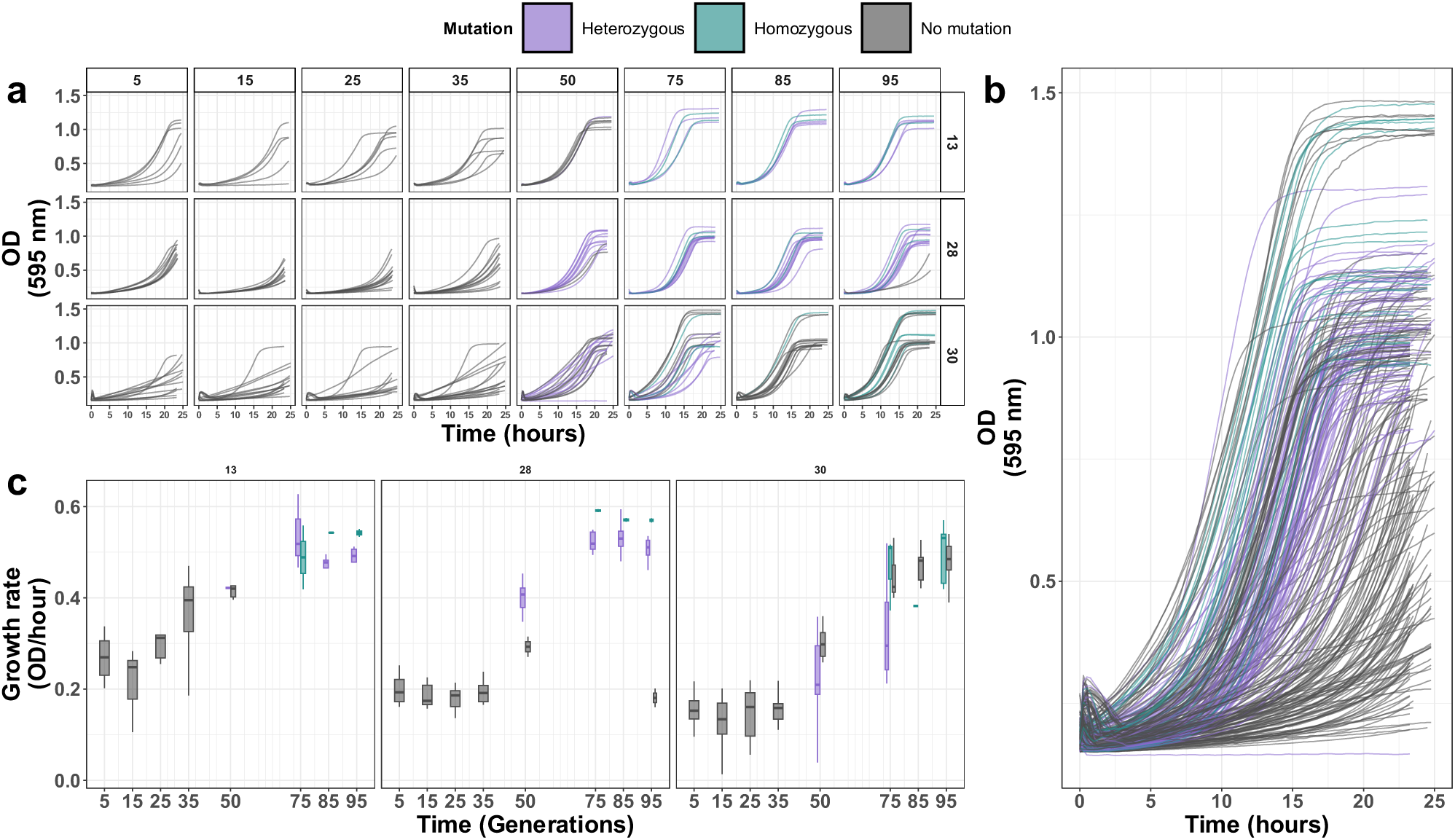
Fitness dynamics of *PDR1* mutants across hybrid lines in which an LOH was detected (Fig. 3b). **a,** Optical density as a function of time in UV mimetic conditions (4 µM of 4-NQO) of single colonies isolated through time (5 ,15, 25,35,50,75,85, and 95 generations) in different lines (13, 28, and 30) (*n* = 270). **b,** Optical density as a function of time (measured in hours) in UV mimetic conditions (4 µM of 4-NQO) across all generations and lines. A pattern emerges, where homozygous lines exhibit superior growth compared to heterozygous or lines with ancestral WT sequence. **c,** Growth rate as a function of time (measured in generations) in UV mimetic conditions (4 µM of 4-NQO) for each line (13, 28, and 30) (*n* = 270). Colors across the figure represent different genotypes detected for the *PDR1* locus: heterozygous (*PDR1* mutation present on only one chromosome), homozygous (resulting from an LOH event where both chromosomes have the same *PDR1* mutation), and ancestral WT sequence (no *PDR1* mutation detected).

**Supplementary Fig. 1.**
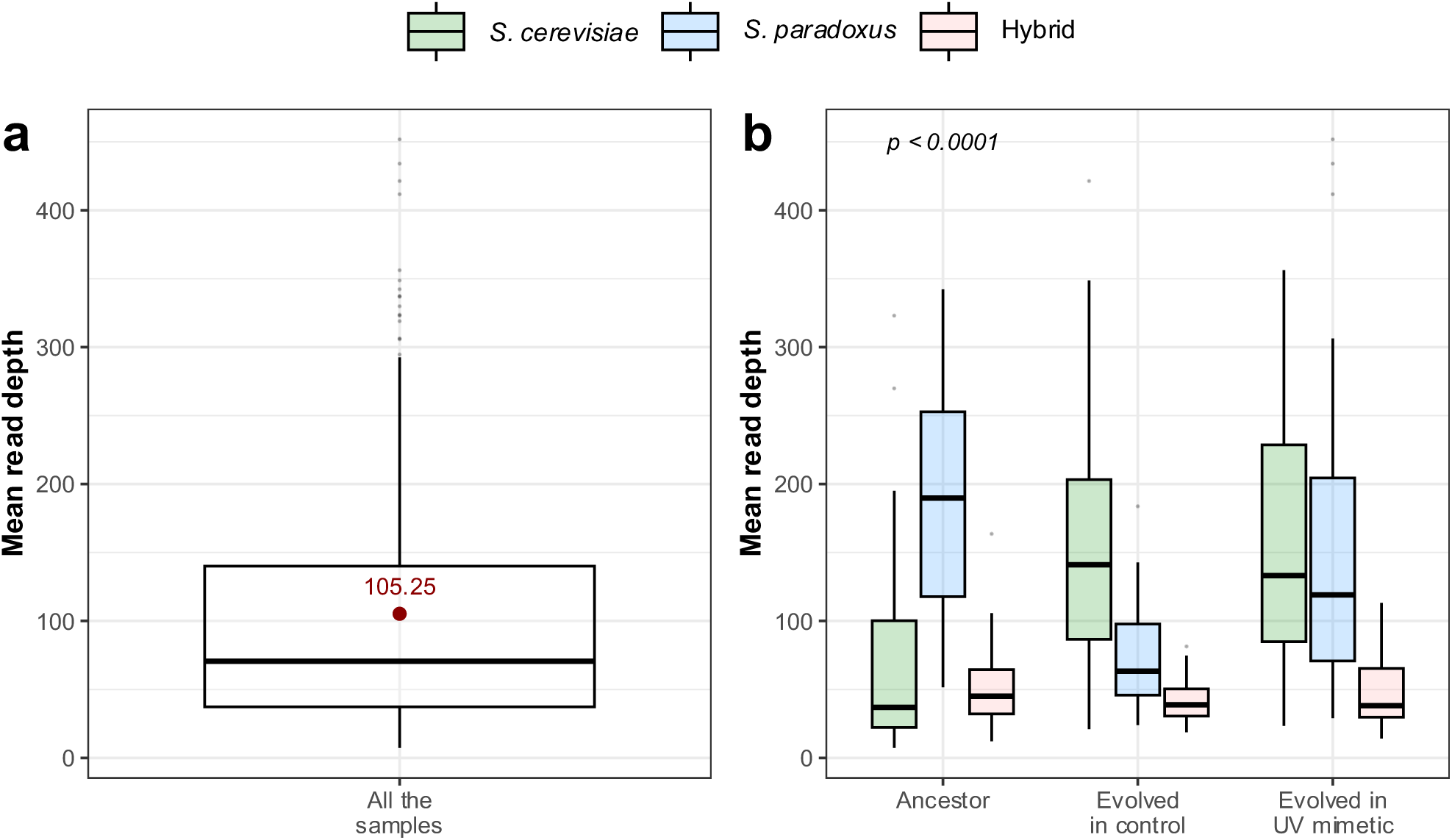
Whole Genome Sequencing (WGS) metrics. **a**, Mean read depth for all sequenced lines (*n* = 300 lines). Mean value is shown in red. **b**, Mean read depth is shown for *S. cerevisiae*, *S. paradoxus* and Hybrid; for Ancestor, evolved in control or UV mimetic conditions (*n* = 30 lines / genotypic background / condition). *p*-value for ANOVA is shown.

**Supplementary Fig. 2.**
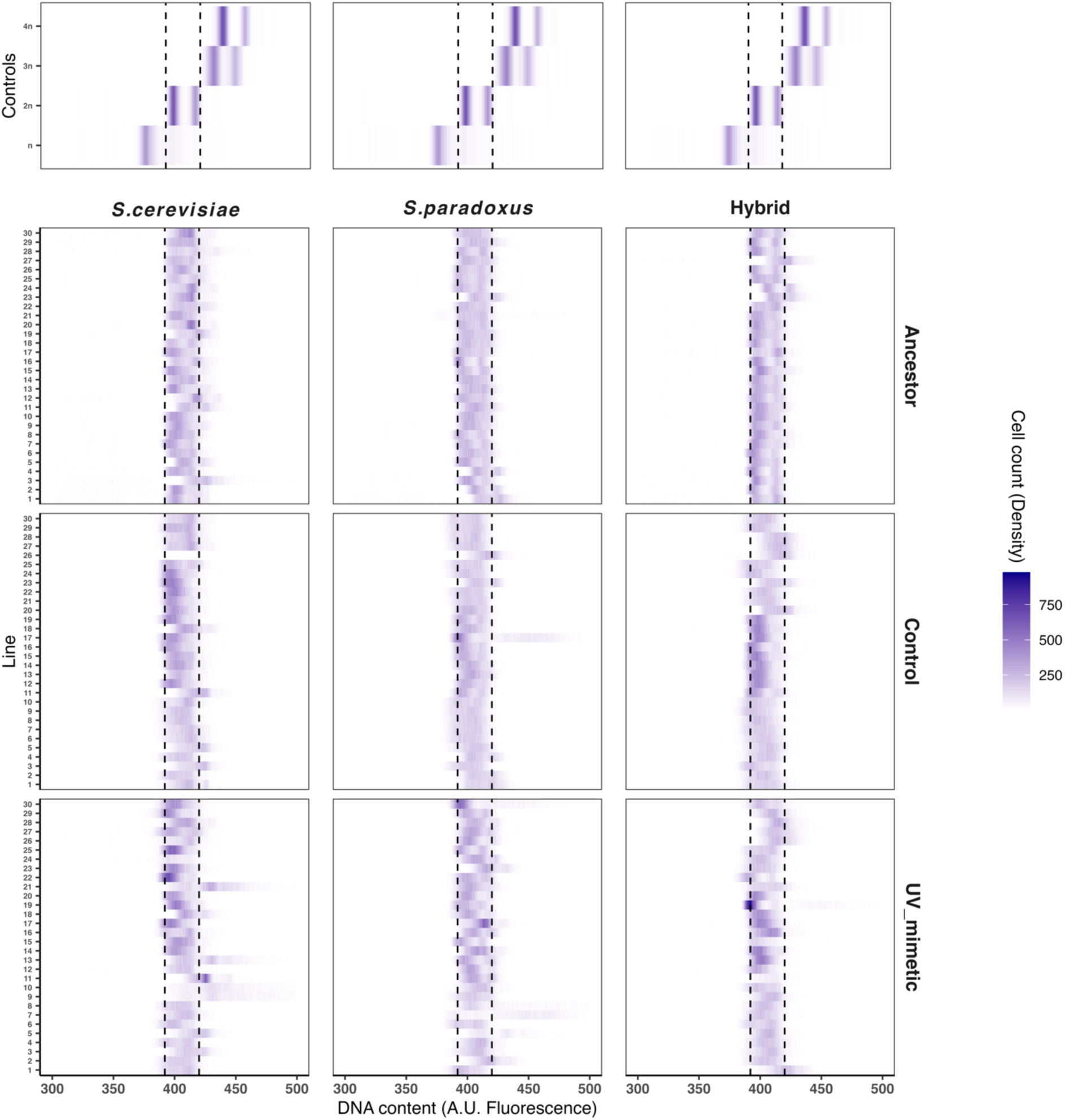
Ploidy dynamics through experimental evolution. DNA content measurements of the 30 lines per genotypic background in ancestral state, control and UV mimetic conditions. DNA content (A.U. Fluorescence) and cell count (density) as gradient color are shown for each line. The dashed line represents 2n states (*n* = 30 lines for each genotypic background and condition). A.U. refers to arbitrary units.

**Supplementary Fig. 3.**
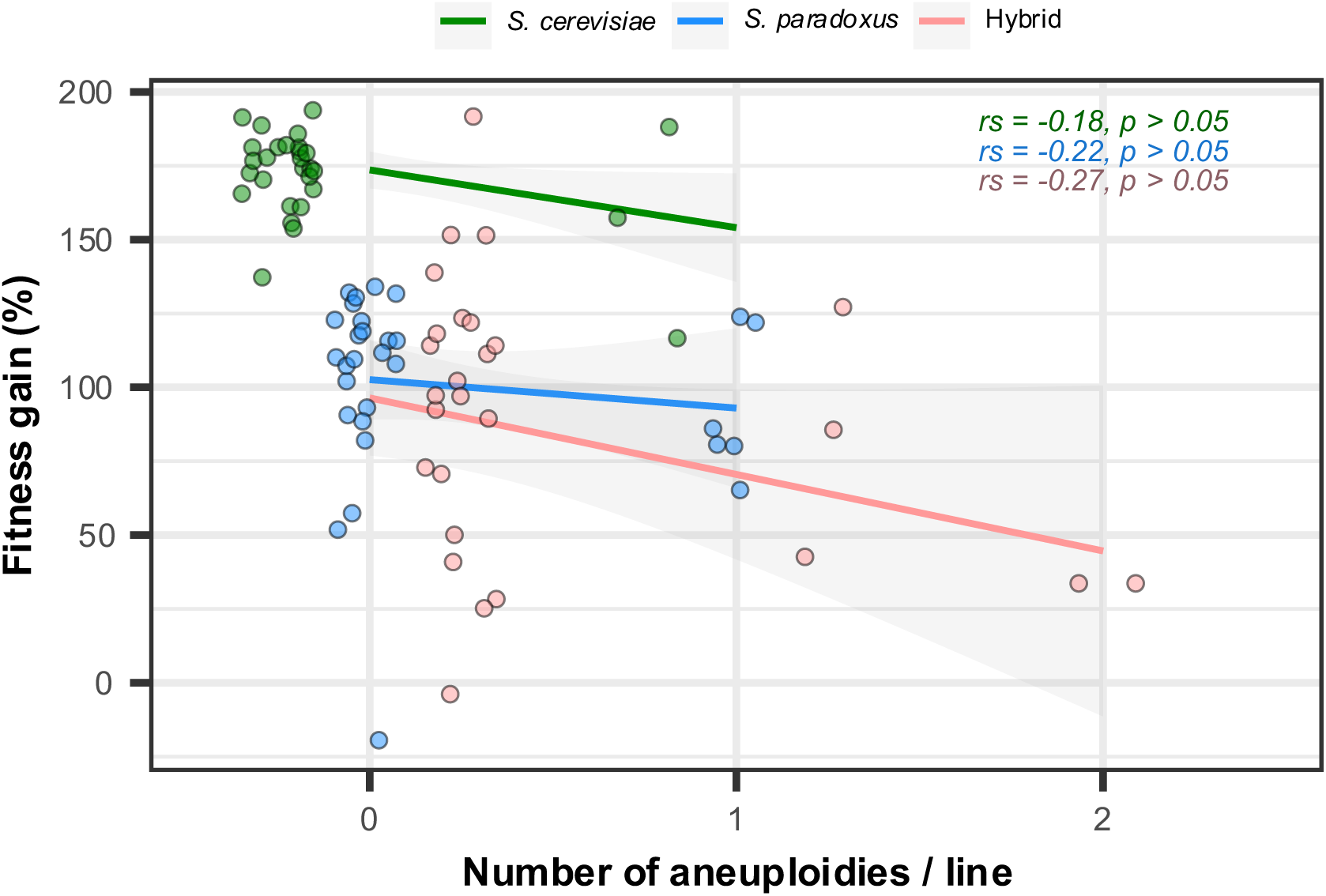
Aneuploidy dynamics. Fitness gain (% change in growth rate between initial and final time points) as a function of the number of aneuploidies/line. Spearman’s rank correlation coefficients (rs) and *p*-values for each genotypic background (*n* = 30 lines for each genotypic background) are shown.

**Supplementary Fig. 4.**
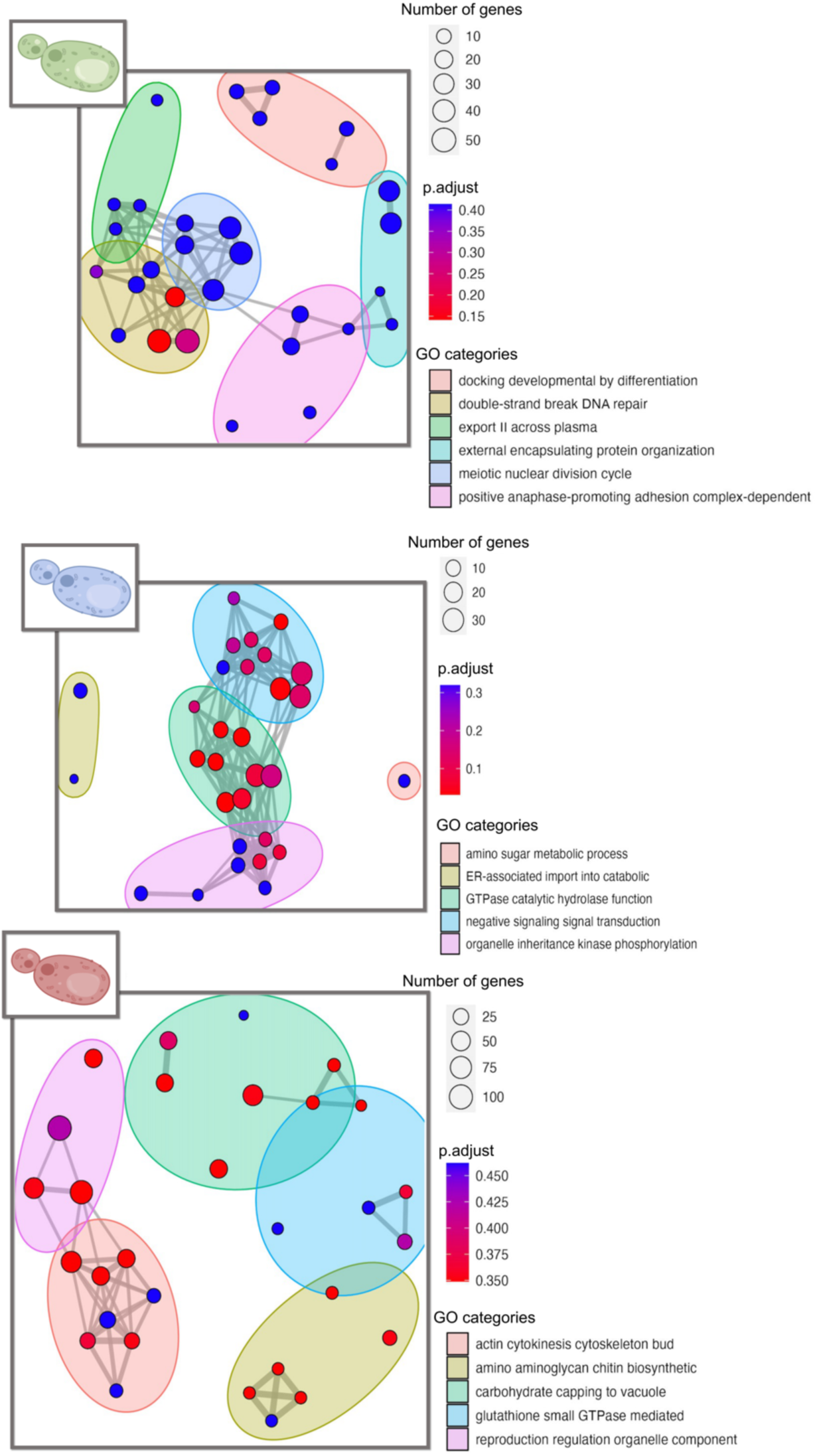
Gene Ontology (GO) enrichment of non-synonymous mutations found in each genotypic background in UV mimetic conditions. Most representative groups are colored for each genotypic background (*S. cerevisiae*, S*. paradoxus,* and hybrid). They are named by *clusterprofiler* function^152^ creating a word cloud of the higher frequency words present in the descriptions of the gene sets included. Each data point symbolizes a distinct GO term, with the associated p-adjust values showcased alongside. False Discovery Rate (FDR) was performed within each genotypic background and adjusted *p*-values are shown (*n* = 765 total non-synonymous mutations for *S. cerevisiae, n* = 270 total non-synonymous mutations for *S. paradoxus,* and *n* = 1326 total non-synonymous mutations for the hybrid). Enrichment ratio was calculated as the foreground fraction to background fraction to detect the most enriched terms: *S. cerevisiae* included double-strand break repair via sister chromatid exchange (GO:1990414) and the regulation of cell differentiation (GO:0045595). *S. paradoxus* included ER-associated misfolded protein catabolic processes (GO:0071712) and endocytic vesicles (GO:0030139). Hybrid comprised trehalose metabolic process (GO:0005991) and ABC-type transporter activity (GO:0140359). Illustrations were created with BioRender.com.

**Supplementary Fig. 5.**
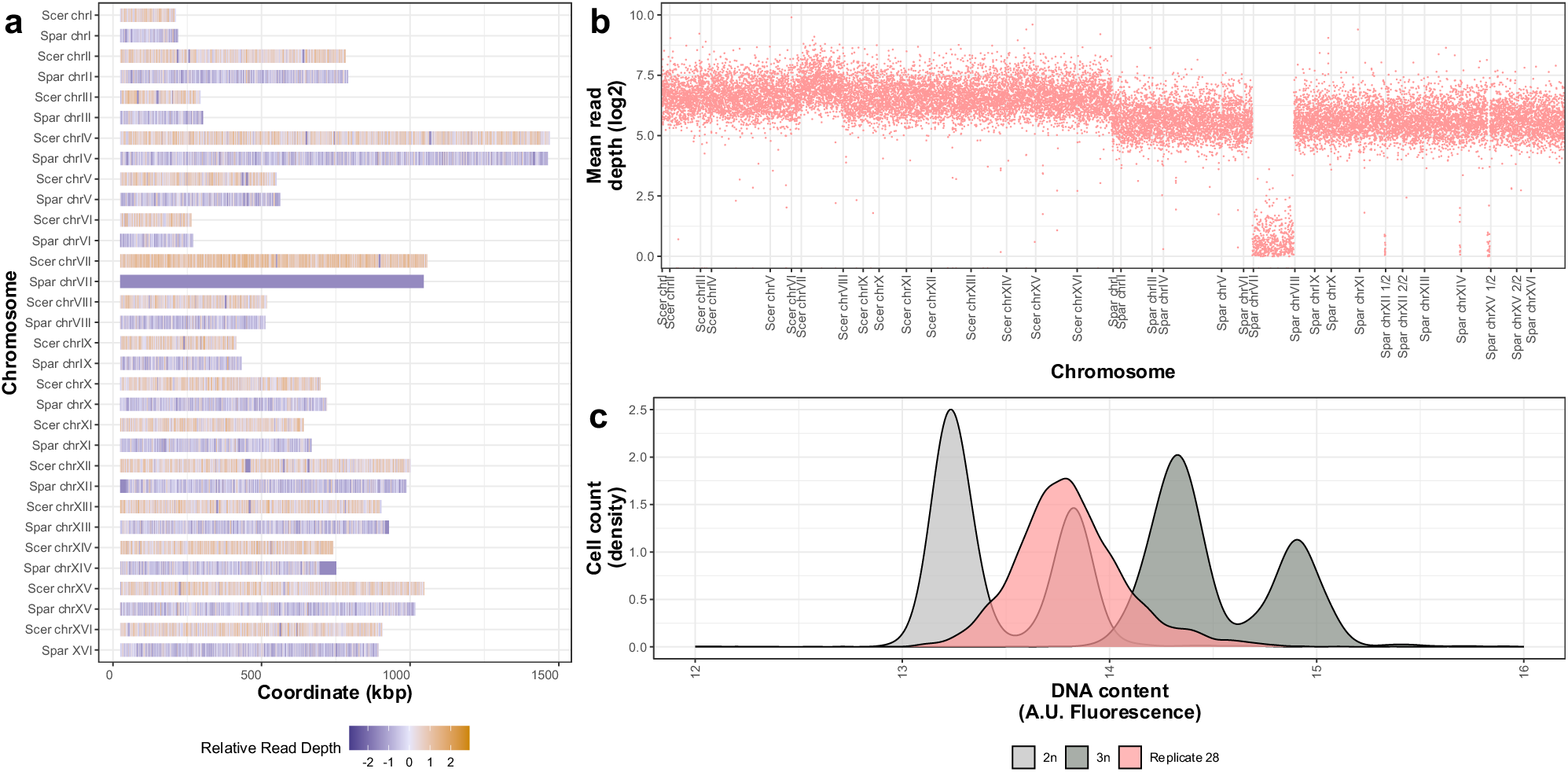
Whole t-LOH in chromosome VII harboring a *PDR1* mutation in hybrid line 28. **a,** Display of relative read depth across chromosomes. **b,** Mean read depth (log2) as a function of chromosome position. **c,** Density of cell count (density) as a function of DNA content (A.U. Fluorescence). A.U. refers to arbitrary units.

**Supplementary Fig. 6.**
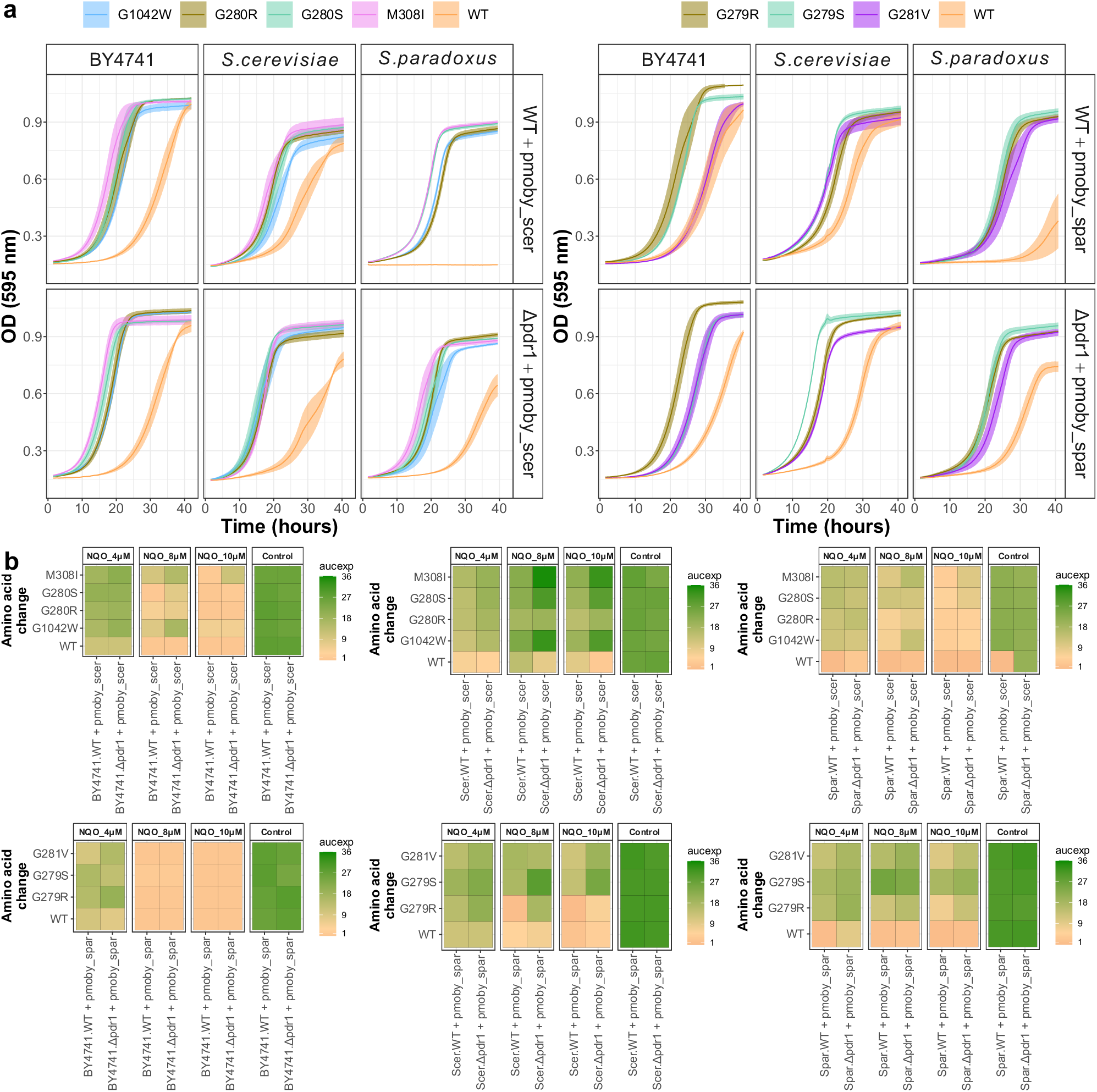
Fitness assay of *PDR1* mutations across genotypic backgrounds. **a,** Optical density as a function of time in UV mimetic conditions (4 µM of 4-NQO) of the *S. cerevisiae* lab strain BY4741 and the natural parental strains (LL13_054 for *S. cerevisiae* and MSH-604 for *S. paradoxus*), all of them, WT (top) or *pdr1*Δ (bottom), expressing *PDR1* variants from a pMoBY plasmid in which the *PDR1* gene is controlled by its native promoter and terminator^156^. The *PDR1* gene was cloned from *S. cerevisiae* background (pmoby_scer) or *S. paradoxus* background (pmoby_spar) (*n* = 4 per growth curve represented with standard error, in total *n* = 216). **b,** Exponential area under curve (AUCexp) across backgrounds (BY4741, *S. cerevisiae,* and *S. paradoxus*) and conditions (4 µM of 4-NQO, 8 µM of 4-NQO, and 10 µM of 4-NQO and control) in both WT or *pdr1*Δ backgrounds expressing *PDR1* mutants from the same plasmids (*n* = 4 per squared, in total *n* = 864).

**Supplementary Fig. 7.**
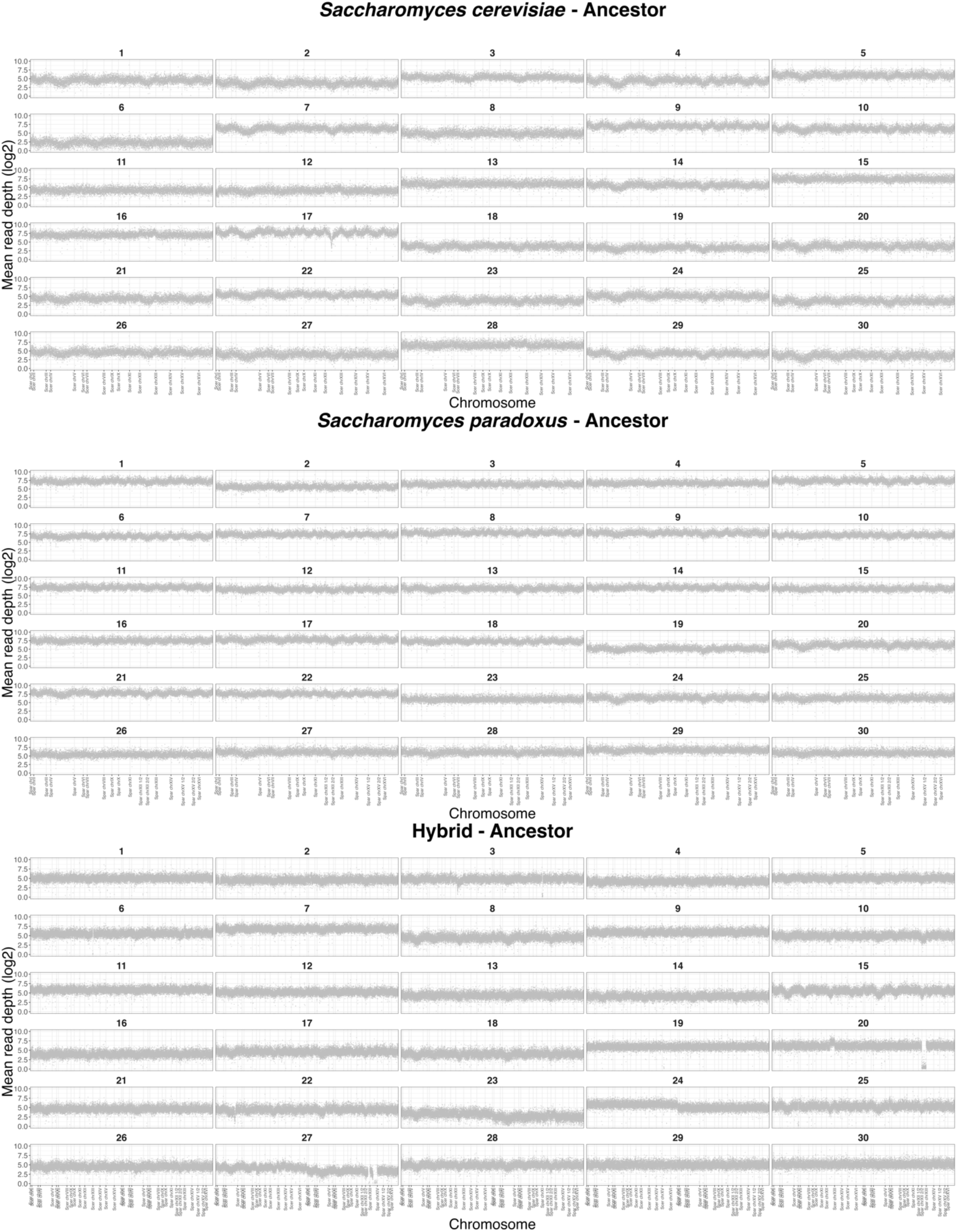
Visualization of average read depth of ancestor lines. Display of mean read depth (log2) across chromosomes (*n* = 30 lines per genotypic background). Chromosomes XII and XV are divided into 1/2 and 2/2 in *S. paradoxus* lines, following the structure of the reference genome (to eliminate repetitive regions). In the hybrid lines, all *S. cerevisiae* chromosomes are positioned on the right, while those of *S. paradoxus* are on the left, facilitating the visualization of ploidy changes (i.e., an increase in one of these complete copies).

**Supplementary Fig. 8.**
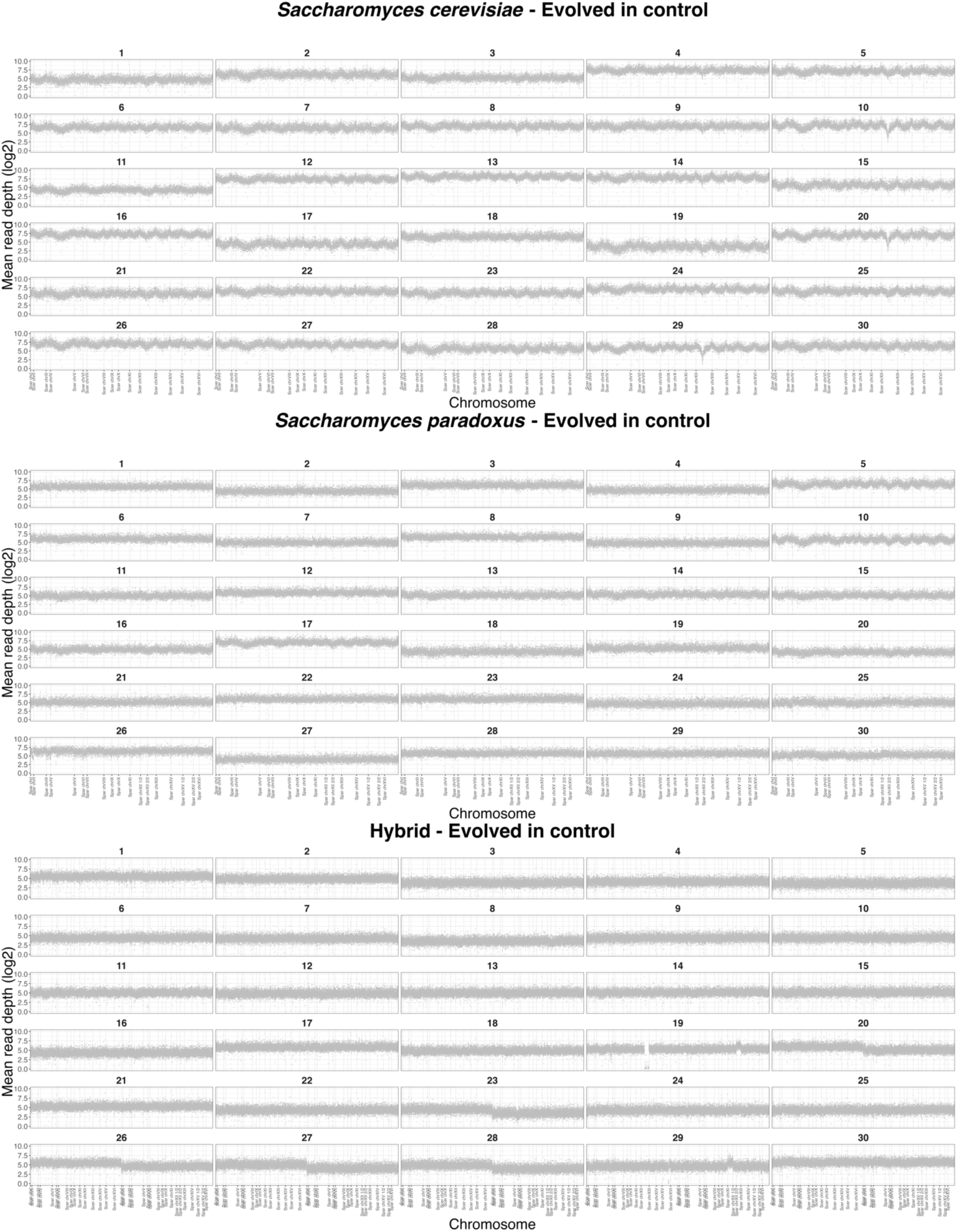
Visualization of average read depth of lines evolved in control conditions. Display of mean read depth (log2) across chromosomes (*n* = 30 lines per genotypic background). Chromosomes XII and XV are divided into 1/2 and 2/2 in *S. paradoxus* lines, following the structure of the reference genome (to eliminate repetitive regions). In the hybrid lines, all *S. cerevisiae* chromosomes are positioned on the right, while those of *S. paradoxus* are on the left, facilitating the visualization of ploidy changes (i.e., an increase in one of these complete copies).

**Supplementary Fig. 9.**
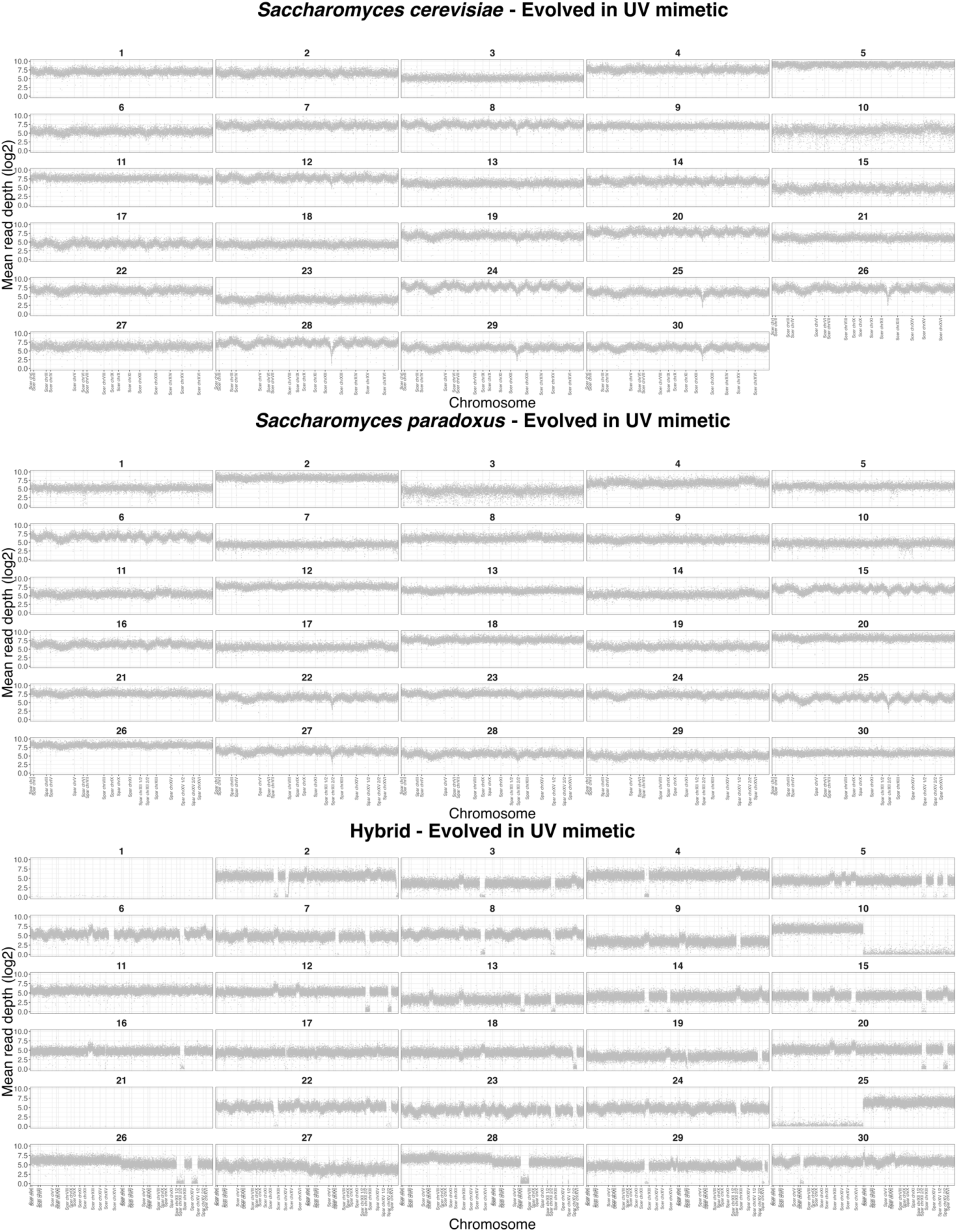
Visualization of average read depth of lines evolved in UV mimetic conditions. Display of mean read depth (log2) across chromosomes (*n* = 30 lines per genotypic background). Chromosomes XII and XV are divided into 1/2 and 2/2 in *S. paradoxus* lines, following the structure of the reference genome (to eliminate repetitive regions). In the hybrid lines, all *S. cerevisiae* chromosomes are positioned on the right, while those of *S. paradoxus* are on the left, facilitating the visualization of ploidy changes (i.e., an increase in one of these complete copies).

**Supplementary Fig. 10.**
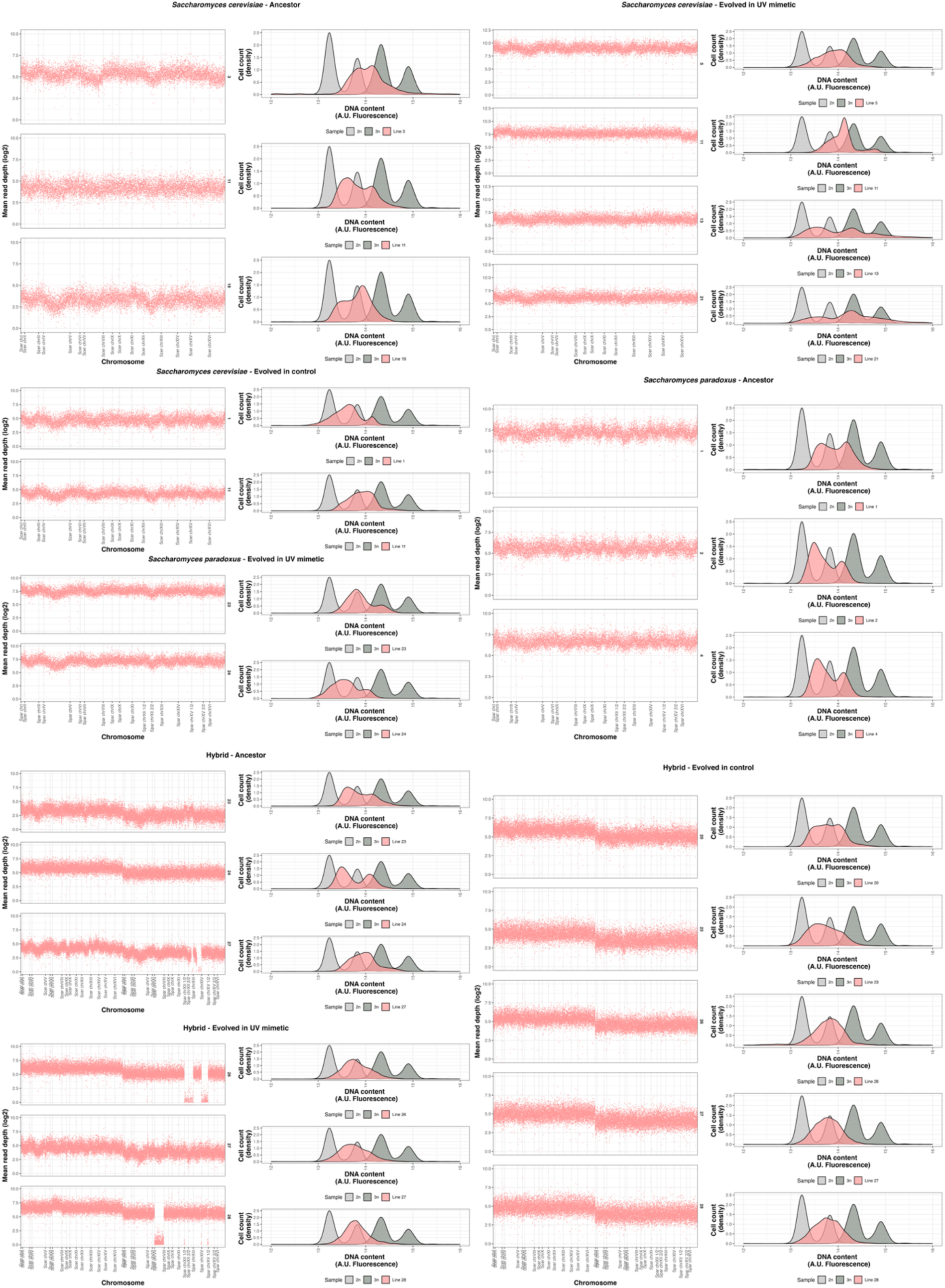
Examples of lines with higher DNA content across conditions and genotypic backgrounds. Mean sequencing read depth (log2) as a function of chromosome position on the left and density of cell count as a function of DNA content (A.U. Fluorescence) on the right (*n* = 3 *S. cerevisiae* ancestor, *n* = 4 *S. cerevisiae* evolved in UV mimetic, *n* = 2 *S. cerevisiae* evolved in control ; *n* = 3 *S. paradoxus* ancestor, *n* = 2 *S. paradoxus* evolved in UV mimetic, and *n* = 3 Hybrid ancestor, *n* = 3 Hybrid evolved in UV mimetic, *n* = 5 Hybrid evolved in control). Individual data points indicate window read depth of ∼1 kbp. 2n and 3n ploidy controls are shown in grey. Chromosomes XII and XV are divided into 1/2 and 2/2 in *S. paradoxus* lines, following the structure of the reference genome (to eliminate repetitive regions). In the hybrid lines, all *S. cerevisiae* chromosomes are positioned on the right, while those of *S. paradoxus* are on the left, facilitating the visualization of ploidy changes (i.e., an increase in one of these complete copies). A.U. refers to arbitrary units.

